# Insula functional connectivity development follows a canonical antero-posterior axis of organization

**DOI:** 10.64898/2026.10.24.761529

**Authors:** Elizabeth A Flook, Golia Shafiei, Matthew Cieslak, Corey Horien, Audrey C Luo, Steven L Meisler, Taylor Salo, Valerie J Sydnor, Tien T Tong, Deanna M Barch, Jeff Goldsmith, Raquel E Gur, Leah H Somerville, Lucina Q Uddin, Aaron Alexander-Bloch, Russell T Shinohara, Theodore D Satterthwaite

**Affiliations:** Department of Child and Adolescent Psychiatry and Behavioral Science, Children’s Hospital of Philadelphia, Philadelphia, Pennsylvania; Penn Lifespan Informatics and Neuroimaging Center, University of Pennsylvania, PA, USA; Department of Psychiatry, Perelman School of Medicine, University of Pennsylvania, PA, USA; Penn-CHOP Lifespan Brain Institute, Perelman School of Medicine, Children’s Hospital of Philadelphia, Philadelphia, PA, USA; Department of Psychiatry, University of Pittsburgh, Pittsburgh, PA, 15213, USA; Department of Psychiatry, Washington University School of Medicine in St. Louis, St. Louis, MO, USA; Department of Psychological & Brain Sciences, Washington University in St. Louis, St. Louis, MO, USA; Department of Biostatistics, Columbia University, New York, NY 10032, United States; Department of Psychology and Center for Brain Science, Harvard University, Cambridge, MA, USA; Semel Institute for Neuroscience and Human Behavior, University of California Los Angeles, Los Angeles, California; Department of Psychiatry and Biobehavioral Sciences, University of California Los Angeles, Los Angeles, California; Neurodevelopment and Psychosis Section, Department of Psychiatry, University of Pennsylvania, Philadelphia; Penn Statistics in Imaging and Visualization Center, University of Pennsylvania, Philadelphia; Department of Biostatistics, Epidemiology, and Informatics, University of Pennsylvania, Philadelphia; Center for AI and Data Science for Integrated Diagnostics (AI2D), Perelman School of Medicine, University of Pennsylvania, Philadelphia, USA

## Abstract

The insula is a heterogeneous region associated with a range of functional domains including sensory, cognitive, and emotional processing. Cytoarchitectural and functional studies have described an antero-posterior axis of insula organization. However, the insula’s functional development remains sparsely described. We delineated the development of insula resting-state functional connectivity across 562 individuals (6-22 years) using generalized additive models and population value decomposition. We show insulo-cortical functional connectivity development differs across the insula: anterior subregions have predominantly decreasing connectivity across the age range examined, whereas posterior subregions have variable connectivity that aligns with the sensorimotor-association axis. Vertex-level analysis revealed that age-related trajectories followed an antero-posterior axis that aligned with estimates of insula microstructure. Our results emphasize that the development of insula connectivity is spatially heterogeneous and aligns with known cytoarchitecture. The normative insula development described here provides a foundation for investigating how emerging psychopathology may be linked to variation in insula development.

## INTRODUCTION

The insular cortex is a heterogeneous brain region supporting a broad range of functional domains, including sensory, cognitive, and emotional processing^1–3^. Located deep to the Sylvan fissure, the insula sits at the confluence of the frontal, temporal, and parietal lobes and has a parallel cytoarchitectural and functional organization. The insula is implicated in multiple large-scale functional networks, including the salience, frontoparietal, cinguloopercular, ventral attention and somatomotor networks^4–8^. The insula is thought to integrate internal and external information and coordinate attentional and affective responses^9^, which are crucial for the affective processing, cognitive control, and social cognition that undergo protracted maturation^10–12^. Despite this, insula functional development in youth remains surprisingly sparsely studied. Here we sought to characterize the development of insula functional connectivity across childhood and adolescence in a large sample.

The insula is organized on an antero-posterior axis, spanning cytoarchitecture, function, and connectivity^13–15^. Consistent with this architecture, structural and functional connectivity vary continuously across the insula from the anterior to posterior pole^13,14^.

Similarly, the cytoarchitecture of the insula is defined by a gradient from granular to agranular cortex ^16–18^ (see **Figure 1**). The posterodorsal insula is granular, meaning it contains densely packed granular cells in cortical layer IV. Gradually, the granular cortex transitions to dysgranular cortex in the middle insula, where granular cells become increasingly less dense. Ultimately, the dysgranular cortex transitions to agranular cortex in the anteroventral insula, where the cortex lacks granular cells.

**Figure 1.**
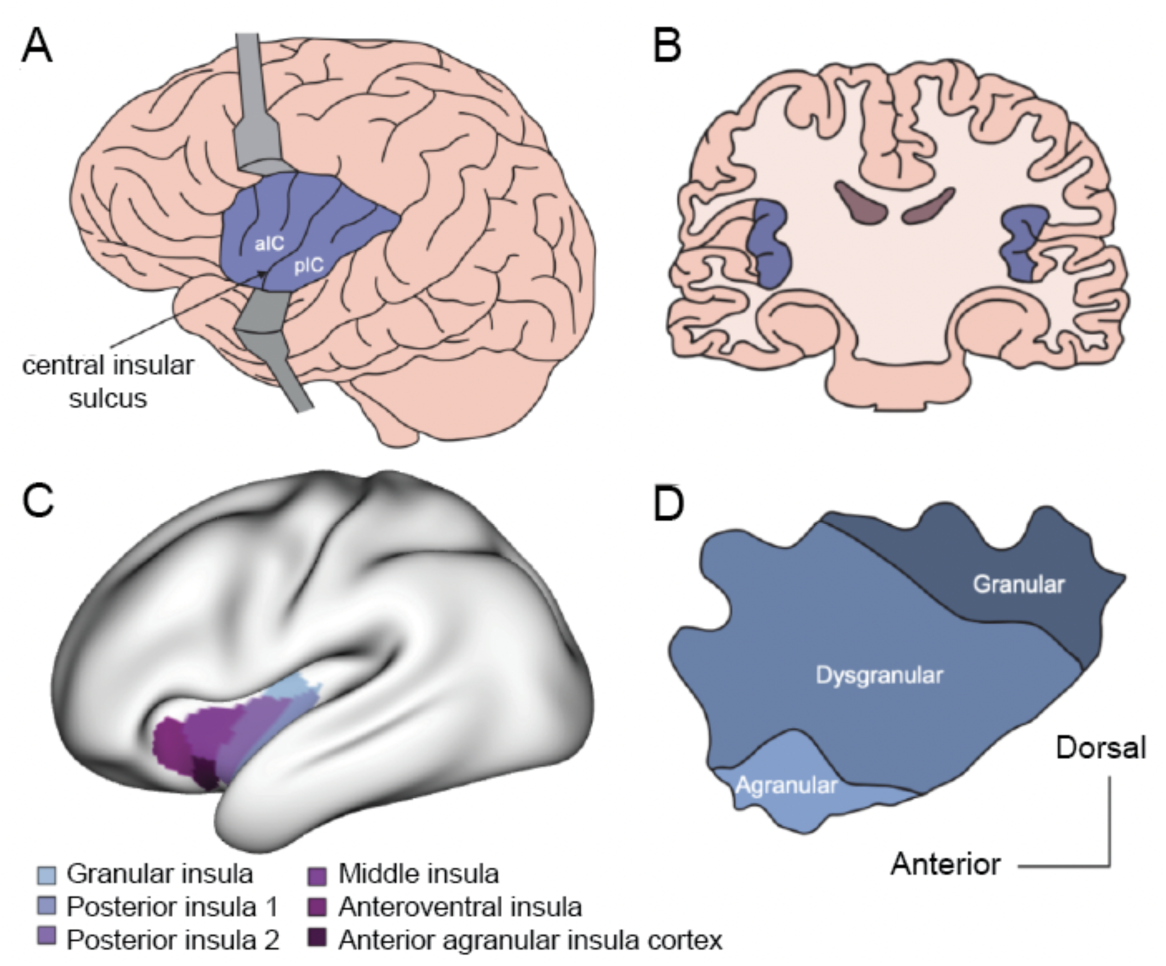
The insula is a region deep to the Sylvian fissure with substantial anatomical and cytoarchitectural heterogeneity. (**a**) the central insular sinus or central insular sulcus provides an anatomical division into anterior or posterior subregions, reproduced with permission from McGregor & LaLumiere 2023 (**b**) the insula is located deep to the confluence of the frontal, temporal, and parietal lobes (**c**) six divisions of the insula as defined in adult data by Glasser et al (2016) informed by function, connectivity, and anatomy (**d**) cytoarchitecture of the insula varies across a gradient, with granular cortex in the posterodorsal insula, agranular cortex in the anteroventral insula, and dysgranular in between.

Functionally, the more posterior regions of the insula are associated with sensory processing, and exhibit strong connectivity with sensorimotor cortices^4,19,20^. In contrast, anterior regions are associated with cognitive and affective processing, and connect with higher-order association networks^4,19–22^. This functional division arises early in insula development; the anterior-posterior distinction can already be seen in children before the age of two^23^.

Despite known insula involvement in cortical functional networks that undergo substantial maturation in adolescence^24–26^, relatively few studies have evaluated the development of insulo-cortical functional connectivity. Whole-brain studies of development often treat the insula as one brain region (for examples see ^26,27^) or only consider a portion of the insula (for examples see ^28–30^). One recent lifespan study focused on the insula demonstrated different age effects on specific functional connections between the anterior, middle and posterior insula subregions across a broad age range (aged 7-54 years old)^31^. This report suggested that insula heterogeneity impacts some age-related trajectories, though these findings were not specific to childhood and adolescence. Structural connectivity studies have likewise demonstrated age-dependent variation in insula connectivity from infancy through young adulthood^32,33^. The few additional studies of insula development in children and adolescents have often focused on group differences between typically developing youth and those with psychopathology, such as autism (for reviews see ^34,35^) and attention deficit hyperactivity disorder (e.g. ^36,37^). Thus, normative age-related trajectories of insula functional connectivity during childhood and adolescence remain surprisingly under-characterized.

Abundant prior work has demonstrated that functional connectivity development is hierarchical and regionally heterogeneous^24,38^. Cortical maturation broadly follows a sensorimotor-to-association hierarchy, with primary sensorimotor systems reaching mature states earlier than higher-order association cortices^39–41^. One framework that integrates these spatial and temporal patterns is the sensorimotor-association axis (S-A axis^42^). The S-A axis organizes brain regions from unimodal sensory cortices to transmodal association cortices. Prior work has shown that functional connectivity development aligns with the S-A axis^43^, with several past studies reporting that connectivity from sensorimotor regions tends to mature earlier than regions in association cortex that undergo protracted development^39–41,44^. The insula spans the S-A axis: subregions within the insula vary from regions processing basic representations of bodily sensations (known as interoception) to regions involved in multisensory integration to regions involved in higher-level processing of emotion and cognition^42,45–49^. As such, the development of the insula’s functional connectivity may be similarly heterogeneous.

In this study, we sought to define insula development across a cross-sectional sample of childhood, adolescence and young adulthood (6–22 years old) to test the hypothesis that development of insula functional connectivity aligns with long-described variation in insula cytoarchitecture and function. Using ultra-high quality resting-state functional connectivity data from the Human Connectome-Development project, we first compared average connectivity in children and adolescents to connectivity in adults.

Next, we identified dominant patterns of age-related changes in insulo-cortical functional connectivity and examined how these patterns compared to patterns of cortical organization such as the sensorimotor-association axis. We integrated subregion-resolved and higher-resolution analyses to test whether age-related trajectories of insulo-cortical functional connectivity aligned with macroscale cortical hierarchies and with known gradients of insula microstructure. As described below, we show that insula age-related trajectories differ along an antero-posterior axis and align with canonical patterns of insular organization.

## RESULTS

We evaluated normative development of insulo-cortical functional connectivity in childhood and adolescence using resting-state functional MRI data from the Human Connectome Project-Development (HCP-D; n = 562, ages 6-22 years) dataset^50^. To characterize overall insula functional connectivity of the developmental sample, we first compared inter-regional insulo-cortical functional connectivity in the HCP-D to an adult sample (Human Connectome Project-Young Adult; HCP-YA, n = 1206, ages 22-35 years)^51^. Next, we investigated the effect of age on insula functional connectivity in the HCP-D sample, characterizing how patterns of development vary across the insula subregions. Finally, we used high-resolution data to evaluate how patterns of insulo-cortical functional connectivity development aligned with insula cytoarchitecture.

### Patterns of insulo-cortical resting state functional connectivity are globally similar in youth and adults

We first defined mean insula resting-state functional connectivity in both HCP-D and HCP-YA. Specifically, we evaluated functional connectivity between the six insula regions and every other cortical region (left hemisphere shown in **Figure 2**, for bilateral results see **Supplementary Figure 1**), as defined by the HCP-MMP1 atlas^52^. In HCP-D, the posterior insula regions were more connected with sensorimotor regions, whereas anterior insula regions were more connected with regions in the association cortex. To directly test how average insula functional connectivity in children and adolescents compared to that seen in adults, we repeated this procedure using the data from HCP-YA and statistically compared the results. We found that insula functional connectivity was quite similar in HCP-D and HCP-YA (correlations for all subregions > 0.91, FDR *p*spin < 0.001), suggesting that global patterns of insula resting-state functional connectivity are already established before adulthood.

**Figure 2.**
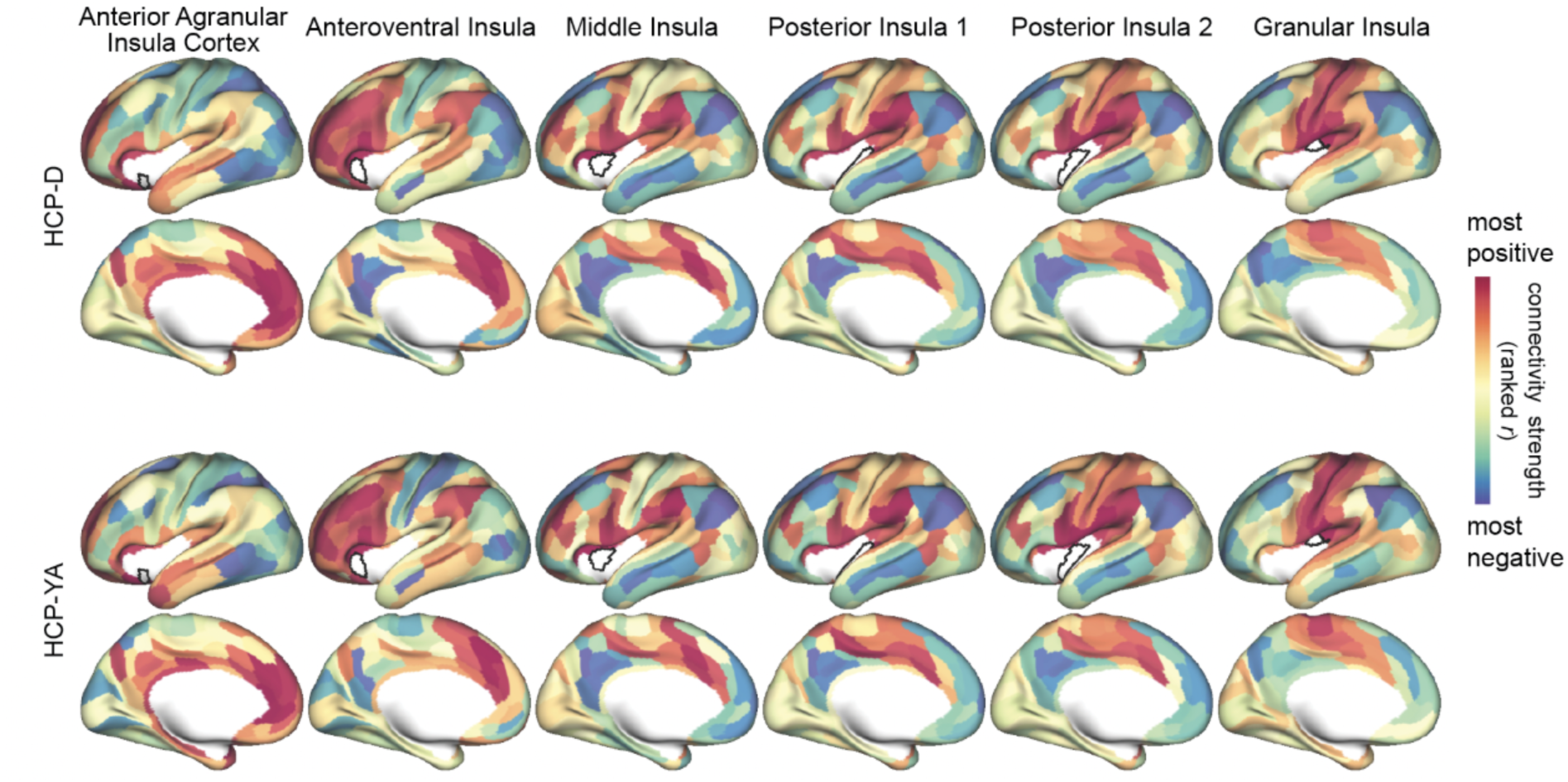
Connectivity to the cortex varies across insula subregions and is globally similar between youth and adulthood. Cortical surface maps rank order of insulo-cortical connectivity between HCP-D (top two rows) and HCP-YA (bottom two rows) for each insula subregion. Subregions are organized from most anteroventral on the left (anterior agranular insula cortex) to most posterodorsal on the right (granular insula). Correlations are shown in rank order from strongest positive (red) to strongest negative (blue) connectivity, with minimal connectivity in the middle (yellow).

### Development of functional connectivity differs across insula regions

Next, we sought to describe how functional connectivity of each insulo-cortical edge changes across childhood and adolescence in the HCP-D cohort. To rigorously capture both linear and nonlinear patterns of insula development without overfitting, we used Generalized Additive Models (GAMs)^53^ with penalized splines^54^ to assess the effect of age on insulo-cortical connectivity (i.e., each edge in the insulo-cortical functional connectivity matrix). Age effects were quantified for each insulo-cortical edge as the partial R^2^, calculated by comparing the full GAM model to a reduced GAM model without the age term. Across all 4,176 connections, 45.5% of edges had significant age effects (FDR-corrected *p* < 0.05); these effects included both increases and decreases in functional connectivity with age (left hemisphere shown in **Figure 3**, for bilateral results see **Supplementary Figure 2**).

**Figure 3.**
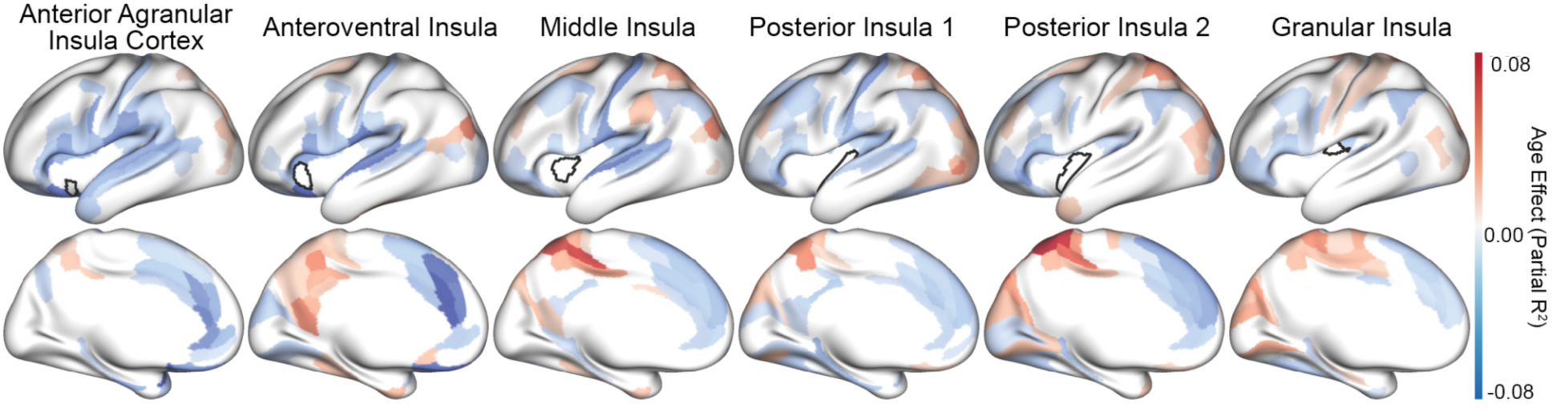
Effect of age on insulo-cortical functional connectivity varies across insula subregions. Cortical distribution of age effects reflects variable patterns of insulo-cortical age-related trajectories. Partial R2 was calculated from Generalized Additive Models (GAM) comparing a model with and without age. Multiple comparisons were accounted for using the false discovery rate (FDR). Colors reflect direction and strength of significant age effects (positive = red, negative = blue, significance defined as p_FDR_ < 0.05), with non-significant subregions in white.

We next sought to characterize the age-related trajectories of the insulo-cortical age effects. We hypothesized that differences across insula subregions in the age-related trajectories of insulo-cortical connectivity would vary with the S-A axis. We reasoned that the S–A axis would provide a meaningful framework for these differences because cortical maturation is organized along this hierarchy, with sensorimotor regions maturing earlier and association regions showing more protracted development^39,41,42^.

Moreover, insula subregions span a broad range of sensorimotor-to-association identities, suggesting that their connectivity may mature differently depending on the position of their cortical targets along the S–A axis. To evaluate the age-related trajectories against the S-A axis, we examined the GAM-derived predicted connectivity across the age range. We visualized the insulo-cortical trajectories by combining cortical regions into quartiles of the S-A axis (**Figure 4a**). The first quartile of the S-A axis encompasses sensorimotor regions. Here, we found that predicted functional connectivity trajectories diverged between anterior and posterior insula subregions: functional connectivity with anterior regions declined across the age range, whereas functional connectivity with posterior regions increased. Conversely, in the fourth quartile of the S-A axis – the association pole of the cortical hierarchy – homogeneous age-related decreases in functional connectivity from all insula subregions were evident.

**Figure 4.**
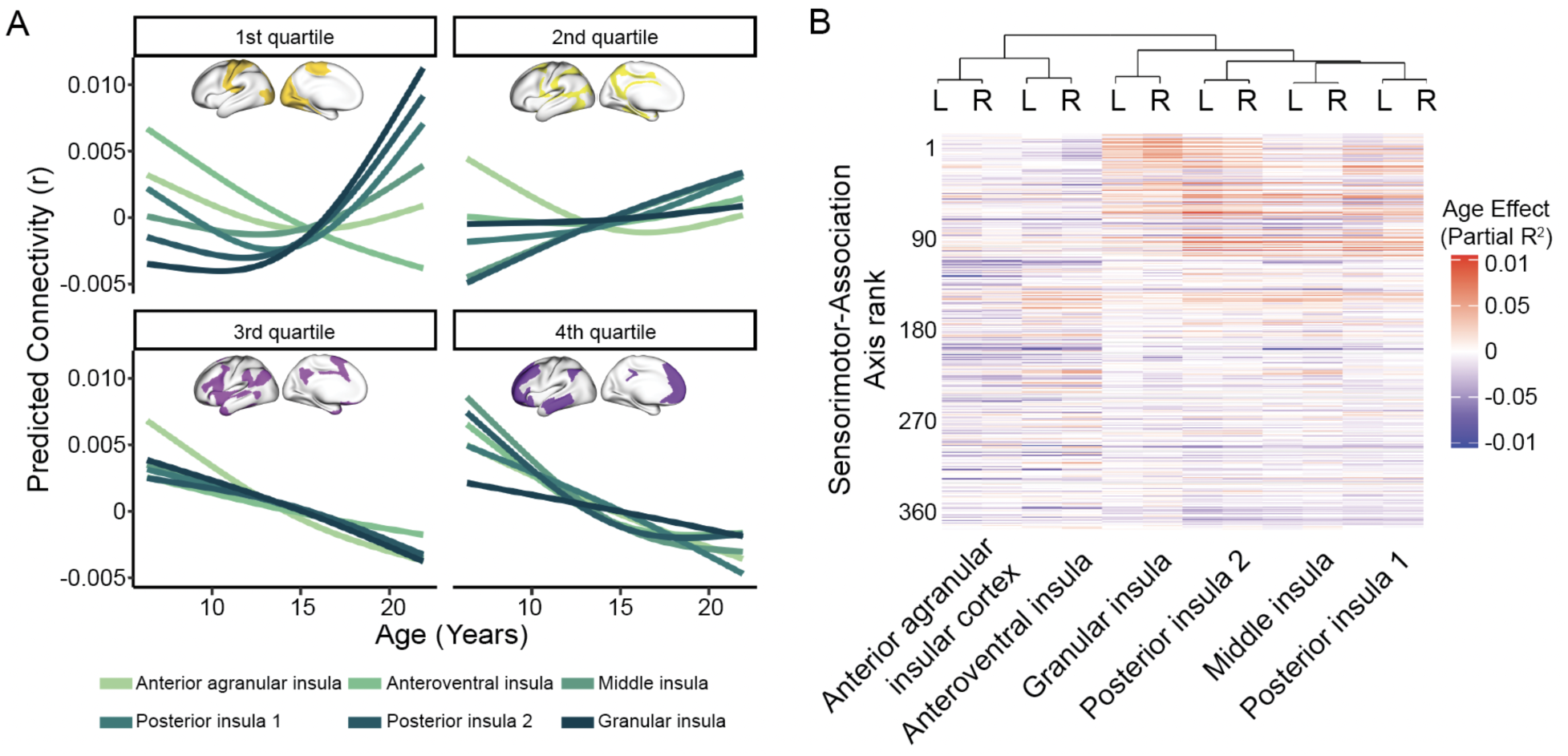
Age-related trajectories of insulo-cortical connectivity from posterior, but not anterior, insula subregions differ across the sensorimotor-association axis. (**a**) For posterior subregions, age-related trajectories of insulo-cortical connectivity varied across the sensorimotor-association axis (S-A axis), whereas the connectivity of anterior insula subregions consistently decreased with age. For visualization, age-related trajectories for each bilateral insula region were averaged within quartiles of the S-A axis defined by the cortical regions. The first quartile represents the sensorimotor pole of the axis; the fourth quartile represents the association pole of the axis (visualized on brain insets). (**b**) Hierarchical clustering of age effects reveals homotopy between bilateral subregions and differentiation between anterior and posterior subregions. Cortical regions (rows) are organized by the S-A axis. Left (L) and right (R) homologous insula subregions clustered together, reflecting parallels between the age-related trajectories of the right and left insula. The highest level of the hierarchy revealed clusters corresponding to bilateral anterior (anteroventral and anterior agranular) versus middle and posterior (middle, posterior 1, posterior 2, and granular) subregions.

To provide a complementary data-driven view of the similarities in the age-related profiles of each insula subregion, we performed a hierarchical clustering analysis of the age effects of insulo-cortical connectivity (**Figure 4b**). The highest level of the hierarchy revealed a split in the age-related profile of anterior and middle/posterior regions. Specifically, regions were clustered into bilateral anterior (anteroventral and anterior agranular) and middle/posterior groups (middle, posterior 1, posterior 2, and granular). Further inspection revealed clustering with the middle/posterior region groups was in part driven by the S-A axis. While the middle/posterior regions appeared to transition from positive to negative age effects across the S-A axis, this was not observed in anterior regions.

In addition to this higher level of organization, the dendrogram’s lowest level revealed that each homologous left and right insula region was clustered most closely together, indicating developmental homotopy. Together, these complementary analyses provided an initial suggestion that development of insula functional connectivity varied along an antero-posterior axis: development of connectivity from posterior insula regions appeared to be more variable and align with the S-A axis, whereas development of anterior insula regions did not.

### Developmental patterns vary along an antero-posterior insula axis

Based on these initial results, as a next step we sought to quantify how patterns of insulo-cortical functional connectivity development varied across insula subregions. To do this, we used population value decomposition (PVD) analysis^55^: a statistical approach that uses two stages of principal components analyses (PCA) to concisely summarize high dimensional patterns in a population (as shown in **Figure 5**).

**Figure 5:**
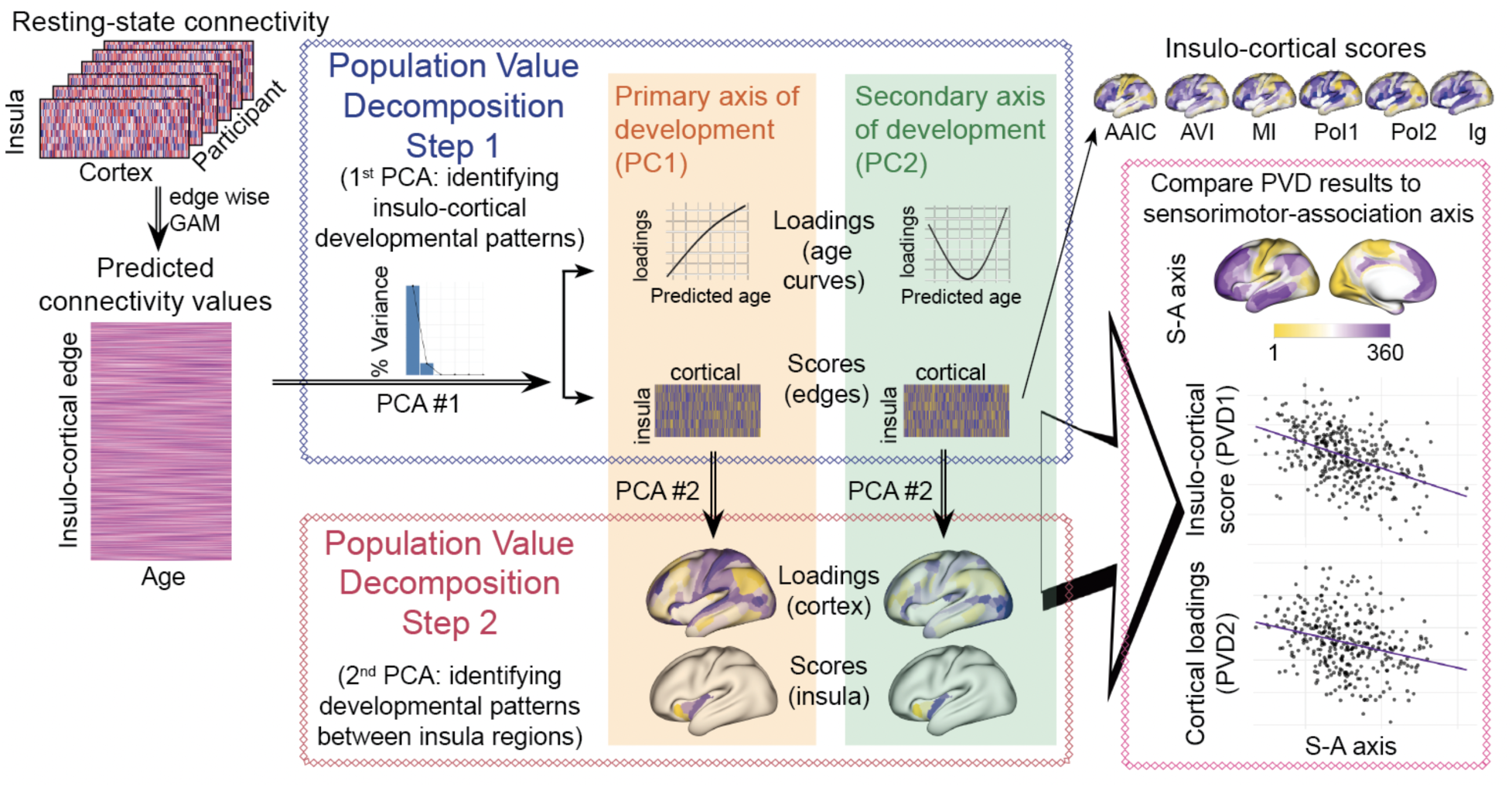
Population value decomposition was used to evaluate cortical patterns of age-related trajectories across insula regions. Schematic describing main analyses, including the population value decomposition (PVD). Resting-state connectivity from participants between the ages of 6 and 22 years were analyzed using Generalized Additive Models (GAMs). GAMs were used to create predicted trajectories of insulo-cortical development from resting state connectivity, which were then submitted to a first principal component analysis (PCA, blue box) to detect the shape of the age-related trajectories (PVD Step 1 loadings; age curves) and the cortical distribution of developmental patterns (PVD Step 1 scores; insulo-cortical edges). The first two principal components accounted for >99% of variance and thus were included in analysis. PCA of age-related trajectories revealed a primary axis of development (PC1: orange shading) and secondary axis of development (PC2: green shading). These developmental scores were then submitted to a second PCA (red box) to detect similarities in patterns of cortical development across insula regions (PVD Step 2, scores). Whole-brain patterns from both step 1 and step 2 were compared to the Sensorimotor-Association axis to detect the underlying structure of the data. AAIC: anterior agranular insula cortex; AVI: anteroventral insula, MI: middle insula; PoI1: posterior insula 1; PoI2: posterior insula 2; Ig: granular insula.

The first step of the PVD answers the question: *is there a dominant pattern shared across the age-related trajectories of insulo-cortical edges?* The input to this first PCA was the modeled age-related trajectory (derived from the GAMs described above) of every connection between each insula subregion and each cortical parcel.

Specifically, this consisted of a matrix of predicted connectivity values for each edge (as rows) and age (as columns). The first component (PC1) of this PCA explained 88.6% of the variance and revealed a predominantly linear pattern of development. The sign and magnitude of each edge’s PC1 score indicate the direction and strength of its expression of this developmental pattern: positive scores reflect greater functional connectivity with age, whereas negative scores reflect lower functional connectivity with age, with scores near zero indicating relatively little age-related change. In contrast, the second component (PC2) explained 11.3% of variance, with a quadratic developmental pattern. For simplicity, moving forward PC1 is referred to as the primary developmental axis, and PC2 is referred to as the secondary developmental axis. For each insula subregion, we visualized the scores of the primary developmental axis across the cortex (left hemisphere results shown in **Figure 6a**, for bilateral see **Supplementary Figure 3**). Anterior insula regions showed predominantly negative PC1 scores, whereas posterior regions exhibited a more heterogeneous mixture of positive and negative scores. Secondary developmental-axis scores are shown in **Supplementary Figure 4**.

**Figure 6.**
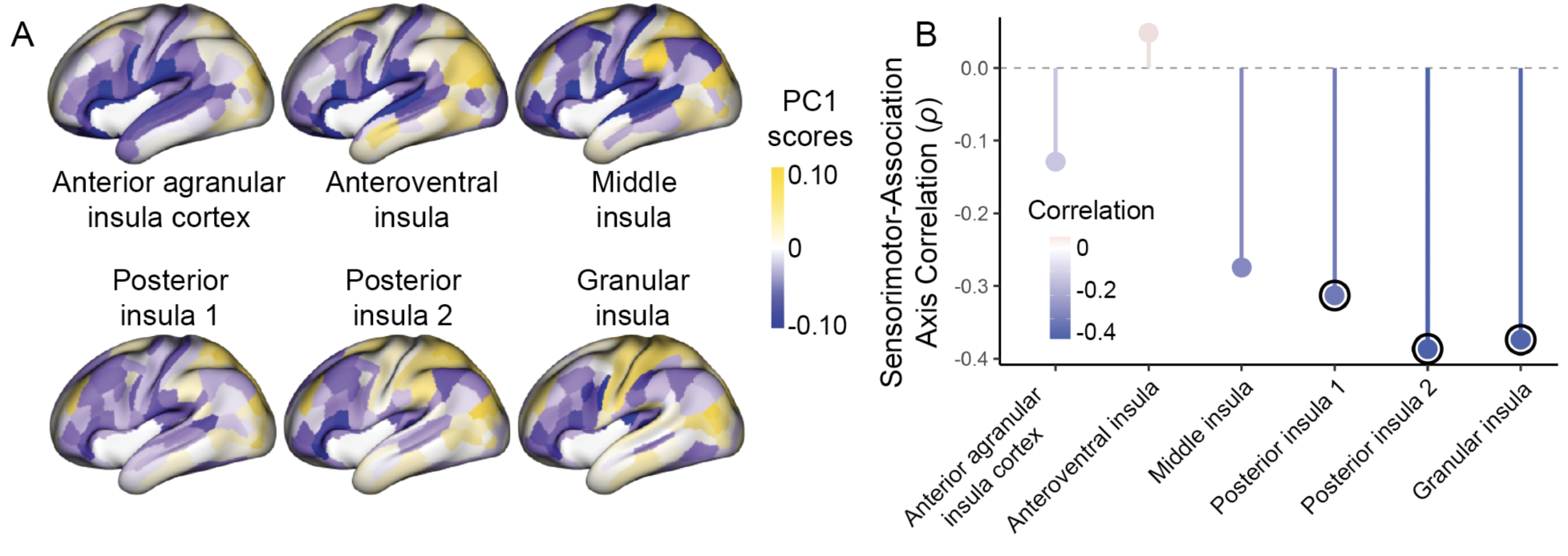
Cortical patterns of insulo-cortical age-related trajectories vary across insula subregions. (**a**) The cortical distribution of PC1 scores along the primary developmental axis of insulo-cortical connectivity varied across insula subregions. These values are the extracted PC1 scores from the first step of the PVD, representing the primary developmental pattern of connectivity across all edges, arranged such that the individual insula regions are visualized separately. Positive scores reflect greater connectivity at older ages; negative scores reflect lesser connectivity at older age. (**b**) Correlation between the PC1 scores and the sensorimotor-association axis (S-A axis) demonstrated that the alignment between S-A axis and development patterns is most pronounced in the posterior regions of the insula. Positive correlation shown in red, negative correlation shown in blue. Significant correlations (p_spin_ < 0.05) circled in black.

We next sought to descriptively annotate the primary developmental axis of insulo-cortical connectivity for each insula subregion, given the heterogeneous developmental patterns visualized between different insula subregions. This heterogeneity was most notable between anterior and posterior regions. As suggested by our initial clustering analysis, we hypothesized that the overall development of insulo-cortical functional connectivity would unfold across the S-A axis. Accordingly, we examined the association between the scores of the primary developmental axis and the S-A axis for each insula subregion using Spearman correlations **(Figure 6b)**. The cresults varied across insula subregions, demonstrating alignment with the S-A axis in the posterior insula subregions that was not seen in the anterior subregions.

As the primary developmental axis aligned with the S-A axis in the posterior insula only, we next examined if insulo-cortical developmental patterns in the anterior insula aligned with other major features of brain organization – like canonical functional networks. To do this, we pursued a network enrichment analysis in the anterior insula using the functional networks defined by Ji and colleagues^56^. We also evaluated network enrichment for the most posterior insula subregion (granular) as a comparison, to determine if network enrichment findings were selective for the anterior regions. For each of these insula regions, we compared the average score for each cortical network to a conservative, autocorrelation-preserving null distribution (see methods). This procedure revealed the agranular insula’s cortical pattern of development was enriched for the cingulo-opercular (*pspin* = 0.006), language (*pspin* = 0.035), and the orbitoaffective (*pspin* = 0.037) networks. The anteroventral insula’s cortical pattern of development was enriched for the dorsal attention (*pspin* = 0.001), cingulo-opercular (*pspin* = 0.010), and default mode (*pspin* = 0.032) networks. Only anteroventral-dorsal attention survived multiple comparison correction (*p*FDR = 0.036) The granular insula did not demonstrate enrichment with any networks (for full results see **Supplementary Table 1**).

### Insula development is organized along an antero-posterior gradient that reflects differential alignment with the sensorimotor-association axis

Our initial results provide insight into developmental patterns of insulo-cortical edge connectivity. We next sought to answer the question: *how is development spatially organized across the insula?* To capture this variation systematically, we continued to the second step of the PVD, which applies a second PCA to the results of the first PCA. This second stage PCA was applied separately to the scores of the primary developmental axis and the scores of the secondary developmental axis (e.g., the first and second components of the first stage PCA). This allowed us to move from an edge-centric view of development (as seen in **Figure 6**) to a regionally-focused view, comparing scores for each insula subregion and loadings for each cortical parcel. This procedure revealed a clear gradient of differential development between anterior insula subregions compared to the posterior subregions (left hemisphere results shown in **Figure 7a**, for bilateral results see **Supplementary Figure 5a)**. This antero-posterior ordering was also seen for the secondary developmental axis (**Supplementary Figure 6a)**, emphasizing that resting-state insula connectivity development occurs along an antero-posterior gradient.

**Figure 7.**
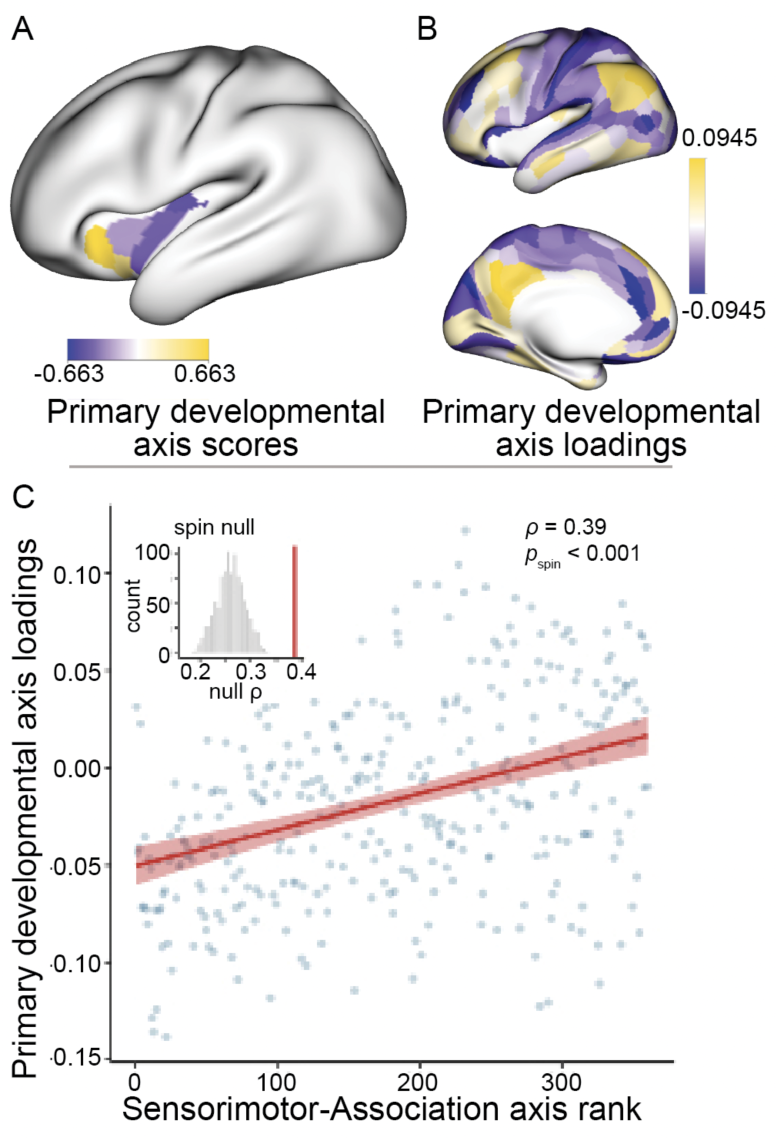
Summary of the primary development pattern demonstrated heterogeneity across the insula and cortex. (**a**) The patterns of primary insulo-cortical development reflected an insula distribution such that the anterior regions had a similar pattern of cortical development that differed from the pattern seen in posterior regions. Scores were extracted from the second step of a population value decomposition (PVD), which used two PCAs to extract patterns of insulo-cortical development from predicted connectivity across children, adolescents, and young adults. (**b**) Cortical loadings of primary developmental patterns extracted from PVD. (**c**) Cortical loadings from the primary developmental pattern have a positive linear association with the S-A axis, with spin tests showing a significant correlation between cortical loadings and S-A axis as compared to a null distribution (inset). Red shaded areas represent 95% confidence intervals around the fitted mean.

To better understand what pattern of cortical connectivity was represented by the antero-posterior gradient across the insula, we evaluated the cortical loadings from the second stage of the PVD. Based on our results thus far, we expected that this pattern would align with the S-A axis. Notably, the cortical loadings from the primary axis of development showed a positive association with the S-A axis (*r* = 0.39, *pspin* < 0.001; left hemisphere results shown in **Figure 7b**, for bilateral results see **Supplementary Figure 5b**); this was also seen for the secondary axis of development (*r* = 0.39, *pspin* < 0.001; **Supplementary Figure 6b**). Taken together, these results emphasize that insula development is organized along an antero-posterior gradient that reflects differential alignment with the S-A axis.

### Development of the insula functional connectivity aligns with insula microstructure

Our results thus far provide evidence that insulo-cortical functional connectivity undergoes heterogeneous development, with a gradient from posterior to anterior regions. Notably, the insula has a well-described cytoarchitectural gradient along the antero-posterior axis, in which granular cortex in the posterodorsal insula transitions to dysgranular cortex in the middle insula and agranular cortex in the anteroventral insula^3,18,57^. As a final step, we sought to test the question: *does the insulo-cortical functional connectivity development align with the insula’s underlying cytoarchitectural gradient?* Answering this question required a higher-resolution approach that could capture fine-grained insula variability. We used a vertex-level approach, evaluating functional connectivity between insula and cortical vertices. As our prior results had revealed that insulo-cortical connectivity development was largely explained by linear and quadratic forms, for computational efficiency we modeled this high-resolution data using polynomial regression. Next, we extracted linear and quadratic age coefficients and conducted a PCA to generate a vertex-level map of insula development. This map was then compared to the microstructural brain maps from BigBrain (called MicroG*)^58^ as a validated proxy for cytoarchitecture^59^. We extracted the primary microstructural scores from the insula to compare vertex-level patterns of development from across the insula to the underlying microstructure (left hemisphere results visualized in **Figure 8a**, for bilateral results see **Supplementary Figure 7a**). To assess the significance of the association, we used Moran randomization^60^ to generate a null distribution of correlations while preserving the spatial autocorrelation within the insula. We found that the microstructure was strongly associated with developmental patterns across the insula (left insula and microstructural gradient *r* = 0.60, *pMoran* = 0.015, **Figure 8b**; right insula and microstructural gradient *r* = 0.66, *pMoran* = 0.003, **Supplementary Figure 7b**). Our findings emphasize the development of insulo-cortical functional connectivity closely aligns with the underlying cytoarchitecture.

**Figure 8.**
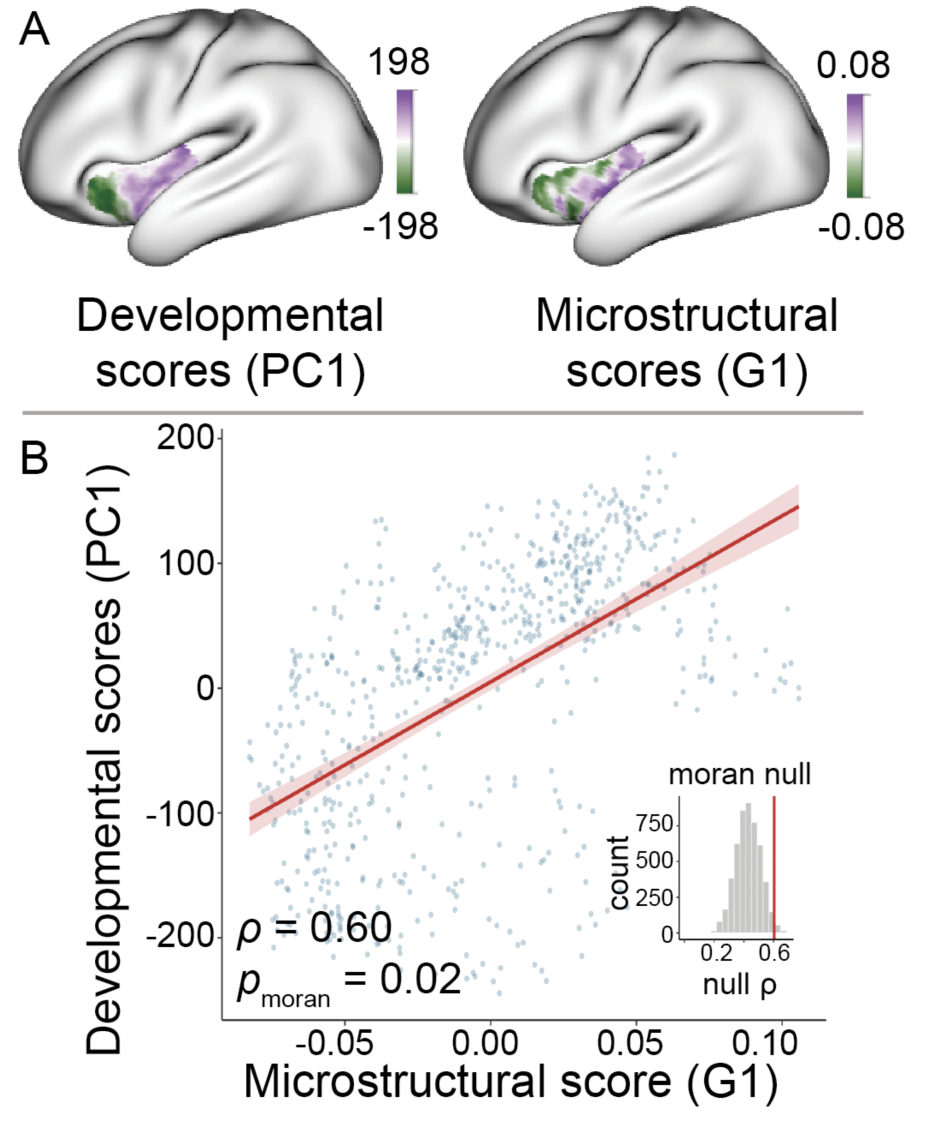
Developmental gradient of the insula aligns with underlying microstructure of the insula. (**a**) Vertex-level analyses demonstrate a transition from anterior to posterior insula, reflecting a similar pattern of age-related trajectories as the parcellated approach. Values reflect the first principal component of insulo-cortical development as defined by linear and quadratic age coefficients. The left figure shows the first principal component, representing the dominant developmental pattern of vertex-level analysis. The right figure represents the dominant underlying microstructural gradient as determined by diffusion map embedding from Paquola et al., 2019. (**b**) Developmental patterns across the insula correlate with underlying microstructure (left insula correlation with primary microstructural gradient shown; ρ = 0.60, p_Moran_ = 0.02). Correlation values determined by Spearman correlation; statistical significance evaluated using Moran randomization.

### Minimal age-related changes in insulo-subcortical functional connectivity

While our effort was focused on the development of insulo-cortical connectivity, we also examined subcortical connectivity. Subcortical regions were defined by the Scale I Tian atlas^61^, which parcellates the subcortex into 16 regions: left and right hippocampus, anterior thalamus, posterior thalamus, putamen, caudate, amygdala, nucleus accumbens, and globus pallidus. Functional connectivity was extracted such that each individual had an averaged connectivity value calculated for each bilateral insula region and subcortical region pair. Development was evaluated using GAMs for insulo-subcortical connections. After correcting for multiple comparisons, only one insulo-subcortical connection showed a significant age effect: the middle insula with the globus pallidus (*p* = 0.037, for complete results see **Supplementary Table 2**).

## DISCUSSION

We show that the canonical antero-posterior organization of the insula also explains the development of insulo-cortical functional connectivity in youth. Functional connectivity from anterior insula regions to the cortex declined with age, whereas development of posterior insula connectivity was more variable and aligned with the S-A axis. Developmental patterns of insula functional connectivity strongly cohered with known cytoarchitecture. Together, these results emphasize that the development of insula functional connectivity should be understood in the context of fundamental properties of insula organization.

The insula’s cytoarchitectural gradient is paralleled by a well-established gradient of functional organization in adults. Specifically, the posterior insula is most associated with sensory processing, the middle insula with polymodal integration, and the anterior insula with cognitive and limbic function (for reviews see ^1,2,9,62,63^). Furthermore, multimodal functional and structural connectivity studies in adults show insula connectivity parallels function, such that the posterior insula has the greatest connectivity with sensorimotor regions and the anterior insula has greatest cognitive and limbic connectivity (e.g. ^4,20,21,64^). Our results in children and adolescents cohere with this adult pattern, joining a growing body of literature identifying regional variability in insula connectivity as a core feature of insula organization throughout the lifespan^13,14,23,31^. Prior developmental work reported age-related changes in specific insula connectivity pathways, including decreasing connectivity with subcortical and core interoceptive regions and increasing connectivity with multimodal cortical regions^31^. Our approach builds upon this prior work by delineating the broader spatial organization of developmental connectivity across the insula, revealing how functional connectivity changes are embedded within an overarching developmental axis.

The S-A axis captures a major spatiotemporal pattern of cortical development, linking regional differences in developmental timing across youth to macroscale cortical organization^41,42^. We found the insula’s primary antero-posterior developmental axis is driven by differential alignment with the cortical S-A axis. Specifically, the development of posterior insula connectivity aligned with the S-A axis; this pattern was largely absent in the anterior insula subregions that showed a distributed age-related decline in functional connectivity with the cortex. This distinction is consistent with prior work showing age-related trajectories vary systematically according to position within cortical functional hierarchies^39,41–43^. Similar to the pattern we observed in the posterior insula, cortical regions closer to the sensorimotor pole of the cortical S-A axis exhibit increasing connectivity with sensorimotor cortices and decreasing connectivity with association cortices across adolescence^38,41,43,44^. This pattern is thought to reflect increasing integration within sensorimotor networks alongside segregation of association networks^38,41,44,65^. The strong alignment of posterior insula development with this canonical cortical axis suggests its maturation may follow broader developmental patterns of sensorimotor cortices. In contrast, the absence of alignment in the anterior insula suggests that its age-related trajectory is less well captured by the S-A axis and may instead reflect a different developmental organizational schema. Together these findings suggest the broader insula does not occupy a single position within a cortex-wide developmental hierarchy. Rather, our results indicate that different subregions participate in different large-scale developmental programs, emphasizing the importance of considering the regional heterogeneity of insula subregions in accounts of brain development.

Abundant literature has defined a gradient of cytoarchitecture in the insula that reflects its function. Specifically, the posterior, granular insula with significant thalamic projections has robust sensory processing^66,67^. In contrast, the anterior, agranular insula is marked by layer V cells with long-range projections – von Economo neurons – and has integrative associative functions^68^. One prior neuroimaging study in adults used diffusion MRI to quantify microstructural properties of insula subregions^69^, which closely approximated known cytoarchitectural gradients ^59^. They found insula and anterior cingulate cortex (ACC) subregions with the strongest connectivity showed corresponding microstructural profiles. This finding provides evidence linking local microstructure of insula subregions to large-scale functional networks in which its subregions participate. Here, we provide evidence for a structure-function correspondence in development, showing spatial differences in the development of functional connectivity aligned with underlying insula microstructure. This regional heterogeneity may be traceable to the insula’s cellular origins: there is evidence that distinct insula subregions may arise from distinct progenitor domains and migratory trajectories. A study of fetal diffusion tractography uncovered two distinct tracts from the ventricular zone to the insula in the developing brain^70^. Due to the impediment of the developing basal ganglia, one path curves around the lenticular nuclei to the anterior insula and the other travels from the lateral horn to the posterior insula. This has also been shown in post-mortem embryonic and fetal tissue, with immunohistochemical markers demonstrating insula subregions with distinct developmental origins, migration trajectories, and cytoarchitectural properties before 32 weeks gestational age^71^. These differences persist into adulthood with differential gene expression across the insula^72^. Thus, the divergent age-related trajectories we observed across the insula may represent a systems-level manifestation of cellular specialization established during prenatal cortical development.

These findings should be considered in the context of several limitations. First, as the data used are a cross-sectional sample, the age effects reported are between-participant associations. Future work using longitudinal data could investigate within-individual development variability of the insula. Second, our age window begins at five years old, thus precluding the evaluation of effects that occur in infancy and early life.

Third, our MRI-derived microstructural maps do not directly index the cytoarchitecture of the insula. While the maps used here have been shown to approximate the gradient of the insula’s granular layer, we are unable to directly test the relationship between functional connectivity and cytoarchitecture.

Our results create a framework of insula normative development in childhood and adolescence. The insula participates in multiple large-scale functional systems, including sensorimotor, salience/cingulo-opercular, and control-related networks that undergo protracted development. This timeframe coincides with a high risk period for the emergence of psychopathology^73^. The insula has been implicated in almost all major forms of psychopathology^74^ including addiction, anxiety, mood disorders, schizophrenia and anorexia^34,62,67,75–80^, many of which begin to emerge during adolescence and young adulthood^73^. In this context, an important next step will be understanding how the timing of emerging symptoms of psychopathology interacts with deviations in insula development. In addition, longitudinal data will provide important information about individual variability and the impact of adversity, trauma, and substance use on developmental trajectories of the insula.

## METHODS

We designed this study with standardized, containerized, and publicly available image processing pipelines.

### Participants

We used data from two large-scale datasets: the Human Connectome Project: Development (HCP-D Release 2.0 Release 2.0)^50^ and Human Connectome Project: Young Adult (HCP-YA Release S1200)^51^. We used HCP-D to evaluate patterns of insula functional connectivity development. HCP-YA was used as a comparison of adult participants.

HCP-D (Release 2.0) is a large study of children and adolescents that approximates the demographics of youth in the U.S. Participants were recruited across four sites: University of Minnesota, Harvard University, Washington University in St. Louis, and University of California-Los Angeles. The original sample included n = 652 participants, of whom 631 had available resting-state fMRI data (age range 5-22 years old). All study procedures were approved by a central Institutional Review Board at Washington University in St. Louis.

HCP-YA (Release S1200) is a study of 1206 healthy young adult participants (age range 22-35 years old) collected across Washington University in St. Louis, University of Minnesota, University of Oxford, Saint Louis University, Indiana University, d’Annunzio University, Ernst Strungmann Institute, Warwick University, Radboud University Nijmegen, and University of California at Berkeley. All study procedures were approved by a central Institutional Review Board at Washington University in St. Louis.

In all studies, written informed consent was obtained for all study participants. For participants under the age of 18, written consent was provided by legal guardians and assent from participants. Demographic information for the datasets may be found in Table 1.

**Table 1:** Demographics for HCP-D and HCP-YA final sample.

|  | HCP-D | HCP-YA |
| --- | --- | --- |
| n<sub>final sample</sub> | 562 | 1206 |
| Age range (mean) | 6 - 22 years (14.8) | 22 - 35 years (29.3) |
| Female identifying | 53.4% | 54.5% |
| Self-reported race |  |  |
| *White* | 63.7% | 73.6% |
| *Black* | 11.0% | 16.0% |
| *Asian* | 8.0% | 5.3% |
| *Mixed* | 14.6% | 2.7% |
| *Other/unknown* | 2.7% | 2.5% |

#### Sample selection Age range restriction

The HCP-D dataset was selected due to the inclusion of children and adolescents, the population targeted by our research question. A cohort of healthy young adults (HCP-YA) was used as a comparator group.

#### Medical exclusion

Exclusion criteria included MRI contraindications, medical conditions affecting brain function, and gross neurological abnormalities. In the HCP-D, 21 participants were excluded from the original sample resulting in 610 remaining participants. In the HCP-YA dataset, no individuals were excluded from the sample resulting in 1206 remaining participants.

#### T1w exclusion

Low-quality T1-weighted sMRI images were excluded by the study consortia before processing if they did not survive manual quality assurance. We did not exclude any additional participants.

#### fMRI exclusion

Resting state scans were evaluated for quality using head motion and duration. To identify high-motion outlier fMRI volumes, framewise displacement was calculated using the formula from Power et al.^81^ with a head radius of 50 mm. Volumes with framewise displacement greater than 10.0 mm were flagged as high-motion outliers for the sake of later censoring ^81^. Following postprocessing (below) participants were excluded if all resting-state fMRI scans had mean framewise displacement ≥0.2. For remaining participants, only rest scans with <0.2 mean framewise displacement were included. These rest scans were then concatenated to maximize scan time, which has been shown to increase reliability of functional connectivity^82^. We excluded participants with less than seven minutes of resting-state fMRI data. These exclusions resulted in remaining sample sizes as follows: 562 participants in HCP-D (age range 6-22 years) and 1206 participants in HCP-YA (with no excluded images from the HCP-YA sample).

### MRI acquisition and processing

We used images from the HCP-D and HCP-YA including T1-weighted structural MRI and resting-state fMRI. Image acquisition parameters for both datasets have been previously reported by their corresponding studies^83,84^. Briefly, HCP-D data included 26 minutes of resting-state scans per participant in total, acquired in four runs with the following acquisition parameters: multiband acquisition, 2mm^3^ isotropic voxels, TR=800ms, TE=37ms, flip angle=52°, number of volumes=478. HCP-YA data included 58 minutes of resting-state scans per participant with the following acquisition parameters: multiband acquisition, 2mm^3^ isotropic voxels, TR=720ms, TE=33.1ms, flip angle=52°, number of volumes=1200. We processed images with BIDS apps^85^ using DataLad^86^ to provide a full audit trail and enhance reproducibility. Detailed information on image processing compiled below is derived from methods descriptions generated by each BIDS App, under the CC0 1.0 Universal license. Preprocessing and postprocessing pipelines are as follows:

#### Preprocessing

##### HCP-D Structural data

HCP-D data included in this manuscript were preprocessed using *fMRIPrep* 22.0.2^87,88^, which is based on *Nipype* 1.8.5^85,89^. The T1-weighted (T1w) image was corrected for intensity non-uniformity (INU) with N4BiasFieldCorrection^90^, distributed with ANTs 2.3.3^91^, and used as T1w-reference throughout the workflow. The T1w-reference was then skull-stripped with a *Nipype* implementation of the antsBrainExtraction.sh workflow (from ANTs), using OASIS30ANTs as target template. Brain tissue segmentation of cerebrospinal fluid (CSF), white-matter (WM) and gray-matter (GM) was performed on the brain-extracted T1w using fast (FSL 6.0.5.1:57b01774)^92^. Brain surfaces were reconstructed using recon-all (FreeSurfer 7.2.0)^93^, and the brain mask estimated previously was refined with a custom variation of the method to reconcile ANTs-derived and FreeSurfer-derived segmentations of the cortical gray-matter of Mindboggle^94^. Volume-based spatial normalization to two standard spaces (MNI152NLin6Asym, MNI152NLin2009cAsym) was performed through nonlinear registration with antsRegistration (ANTs 2.3.3), using brain-extracted versions of both T1w reference and the T1w template.

##### HCP-D Functional data preprocessing

For each of the BOLD runs per subject, the following preprocessing was performed using the fMRIPrep pipeline. First, a reference volume and its skull-stripped version were generated by aligning and averaging one single-band reference. Head-motion parameters with respect to the BOLD reference (transformation matrices, and six corresponding rotation and translation parameters) were estimated before any spatiotemporal filtering using mcflirt (FSL 6.0.5)^95^. B0 inhomogeneity maps, or field maps, were identified for each subject and estimated based on two (or more) echo-planar imaging (EPI) references with topup^96^ (FSL 6.0.5.1:57b01774). The estimated *fieldmap* was then aligned with rigid-registration to the target echo-planar imaging (EPI) reference run. The field coefficients were mapped on to the reference EPI using the transform. BOLD runs were slice-time corrected to 0.346s (0.5 of slice acquisition range 0s-0.693s) using 3dTshift from AFNI^97^. The BOLD reference was then co-registered to the T1w reference using bbregister (FreeSurfer) which implements boundary-based registration ^98^. Co-registration was configured with six degrees of freedom. First, a reference volume and its skull-stripped version were generated using a custom methodology of *fMRIPrep*. Several confounding time series were calculated based on the preprocessed BOLD data: framewise displacement (FD) and three region-wise global signals. FD was computed using two formulations following Power (absolute sum of relative motions)^81^ and Jenkinson (relative root mean square displacement between affines)^95^. FD was calculated for each functional run, both using their implementations in *Nipype* (definitions from Power et al., 2014). The three global signals are extracted within the CSF, the WM, and the whole-brain masks. The head-motion estimates calculated in the correction step were also placed within the corresponding confounds file. The confound time series derived from head-motion estimates and global signals were expanded with the inclusion of temporal derivatives and quadratic terms for each ^99^.

The BOLD time series were resampled into standard space, generating a *preprocessed BOLD run in MNI152NLin6Asym space*. First, a reference volume and its skull-stripped version were generated using a custom methodology of *fMRIPrep*. The BOLD time series were resampled onto the following surfaces (FreeSurfer reconstruction nomenclature): *fsaverage*. *Grayordinates* files^83^ containing 91k samples were also generated using the highest resolution fsaverage as intermediate standardized surface space. All resamplings can be performed with *a single interpolation step* by composing all the pertinent transformations (i.e. head-motion transform matrices, susceptibility distortion correction when available, and co-registrations to anatomical and output spaces). Gridded (volumetric) resamplings were performed using antsApplyTransforms (ANTs), configured with Lanczos interpolation to minimize the smoothing effects of other kernels^100^. Non-gridded (surface) resamplings were performed using mri_vol2surf (FreeSurfer).

##### HCP-YA

Processed structural and functional MRI scans were downloaded following the standardized HCP Pipelines, described in detail elsewhere^83^.

#### Postprocessing

##### HCP-D

The eXtensible Connectivity Pipeline-DCAN (XCP-D, version 0.6.0rc6)^99,101^ was used to post-process the outputs of fMRIPrep version 22.0.2^88,102^, RRID:SCR_016216). XCP-D was built with Nipype version 1.8.6^103^ (RRID:SCR_002502). Native-space T1w images were transformed to MNI152NLin6Asym space at 1 mm3 resolution. For each of the BOLD runs per subject, the following post-processing was performed: In order to identify high-motion outlier volumes, framewise displacement was calculated using the formula from Power et al.^81^, with a head radius of 50 mm. Volumes with framewise displacement greater than 10.0 mm were flagged as high-motion outliers for the sake of later censoring ^81^. In total, 36 nuisance regressors were selected from the preprocessing confounds, according to the ‘36P’ strategy. These nuisance regressors included six motion parameters, mean global signal, mean white matter signal, mean cerebrospinal fluid signal with their temporal derivatives, and quadratic expansion of six motion parameters, tissue signals and their temporal derivatives^99,101^. Finally, linear trend and intercept terms were added to the regressors prior to denoising. The BOLD data were converted to NIfTI format, despiked with AFNI’s 3dDespike, and converted back to CIFTI format. Nuisance regressors were regressed from the BOLD data using linear regression, as implemented in Nilearn. Any volumes censored earlier in the workflow were then interpolated in the residual time series produced by the regression. The interpolated timeseries were then band-pass filtered using a second-order Butterworth filter, in order to retain signals between 0.01-0.08 Hz. The filtered, interpolated time series were then re-censored to remove high-motion outlier volumes. The denoised BOLD was then smoothed using Connectome Workbench with a Gaussian kernel (FWHM=6 mm). Processed functional timeseries were extracted from residual BOLD using Connectome Workbench ^104^.

##### HCP-YA

The postprocessing of the HCP-YA data followed an identical pipeline to that described above for HCP-D with the exception of censoring. HCP-D applied censoring whereas HCP-YA did not; this decision was regarding prior studies demonstrating a greater concern for motion in a younger population (such as HCP-D) than young adults^105^.

### Quantification of functional connectivity metrics

To examine developmental changes in insula functional connectivity profiles, we computed functional connectivity from resting-state scans as our primary measure of interest. Functional connectivity was computed by extracting functional time series of BOLD activity for each cortical vertex using Connectome Workbench^106^ and computing Pearson correlations between the time series of each insula vertex and each remaining cortical vertex in R^107^ (version 2025.05.0+496). Insula vertices were identified using HCP-MMP1 atlas^52^ and included all vertices labeled anteroventral insula (AVI), anterior agranular insula cortex (AAIC), middle insula (MI), posterior insula 1 (PoI1), posterior insula 2 (PoI2), and granular insula (Ig). For region of interest analysis, cortical vertices were also parcellated using the HCP-MMP1 atlas and Pearson correlations were averaged within each insula region and cortical region.

### Comparison to adult connectivity

We hypothesized that overall connectivity in children and adolescents would be similar to connectivity in young adults, as many prior studies have demonstrated large-scale networks are established early in development (e.g.^108,109^). To test this, we compared average insulo-cortical connectivity in a well-characterized population of young adults (aged 22-35 years; HCP-YA) with our target population (aged 5-22 years; HCP-D). For each participant in the HCP-YA and HCP-D cohorts, functional connectivity was quantified for all insulo-cortical edge pairs from the regions of the HCP-MMP1 atlas.

Connectivity for each edge was then averaged across participants within each cohort and compared between HCP-D and HCP-YA using Spearman correlation. If connectivity patterns are broadly similar across age groups, this would suggest that large-scale insula organization is already established by early development.

To test whether average connectivity patterns differed significantly between cohorts, we generated null distributions that account for spatial autocorrelation. Because neighboring brain regions exhibit inflated similarity in connectivity values, conventional null models risk false positives. To address this, we employed the spin-based spatial randomization test^110^, or “spin test,” which controls for distance-dependent spatial autocorrelation. For each insula region, we conducted a spin test to compute *p*-values (*pspin*) for the observed insulo-cortical connectivity correlations between HCP-D and HCP-YA.

Spin tests were performed in R^107^, using code adapted from Vasa et al ^111^ (found at https://github.com/frantisekvasa/). The “rotate_parcellation” algorithm generated 1000 new sets of surface coordinates using random spherical rotations. We then used “perm_sphere_p” to generate null distributions by rotating one set of connectivity values based on the new coordinates from rotate_parcellation. We then compared the observed correlation of connectivity values between cohorts to the correlations from the null distribution of 1000 permuted connectivity values, generating a *p*spin. This process was repeated for each insula region.

### Analyses of insulo-cortical connectivity development

The primary goal of this study was to investigate differences insulo-cortical connectivity across the age range of children, adolescents and young adults. To this end, we generated a developmental model to understand the effect of age on connectivity values.

#### Developmental model

Age-related trajectories are frequently nonlinear (e.g. Faghiri et al., 2019). To flexibly capture both linear and non-linear associations between functional connectivity and age, we fit Generalized Additive Models (GAMs) using the mgcv package^112^ (version 1.9-1) in R ^107^. GAMs were fit for each insulo-cortical edge with resting-state connectivity as the outcome, age as a smooth term, and both sex and in-scanner motion as linear covariates:

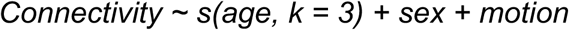

In-scanner head motion was quantified for each participant as mean FD averaged across all included runs. Age was modeled using thin plate regression splines, with the maximum basis complexity (*k*) set to 3 to reduce overfitting. The effect size of age-related changes for each insulo-cortical edge was quantified as the partial R^2^ comparing a full model and reduced model without the age term. The direction of change (increasing vs decreasing connectivity with age) was determined from the average derivative, where a positive derivative indicated increasing connectivity and a negative derivative indicated decreasing connectivity^113^ (version 0.10.0.900). Significant age effects were identified using ANOVA with false discovery rate (FDR) correction (*Q* = 0.05). To visualize estimated smooth function, we generated predicted values from the fitted GAM for 100 evenly spaced values across the age range using *predict* from mgcv package^112^ (version 1.9-1). Age effects were organized into an insula × cortical matrix; Euclidean distances were computed on the transposed matrix using the *dist* function, followed by agglomerative clustering using complete linkage in the *hclust* function from stats package (version 4.4.1). All statistical analyses were conducted in R.

### Developmental patterns of age effects

We used population value decomposition (PVD, **Figure 5**)^55^ analysis to identify consistent patterns of development across the many insulo-cortical connections (inter-insula connections were removed prior to PVD, resulting in a total of 4176 connections). PVD employed a two-step value decomposition. In step one, we applied a principal component analysis (PCA) to the predicted age-related trajectories of insulo-cortical functional connectivity derived from the GAMs. The input matrix consisted of 4176 edges (rows) and 100 ages (columns); PCA was implemented using prcomp (from *stats* package, version 4.4.1). Principal components (PC) 1 and 2 explained the vast majority of variance and were retained for further analysis; PC1 explained 88.6% of developmental variance while PC2 explained 11.3% of the variance. The scores (row weights) of these PCs identified insulo-cortical edges with similar age-related trajectories. The loadings from these PCs (column weights) summarized the shape of the age-related trajectories. For simplicity, PC1 and 2 will be referred to as the primary and secondary developmental axes, respectively.

Next, we sought to understand if these patterns of age-related trajectories differed across insula regions and to investigate if functional properties of the regions were associated with differences in developmental patterns. In step two of the PVD, scores were first rearranged from edge-wise values into two matrices (primary and secondary) of insula regions x cortical regions, which were then each submitted to a second PCA (again using prcomp). For the primary developmental pattern, PC1 explained 48.8% of developmental variance and PC2 explained 23.9%. For the secondary developmental pattern, PC1 explained 43.6% of developmental variance and PC2 explained 23.3%.

The scores from this second PCA represented patterns of development across the cortex for each insula region, allowing for comparison across regions. We also then could use the primary axis of development (PC1 scores) to compare developmental patterns to other metrics of the insula regions.

### Annotation of insulo-cortical edge connectivity development

The results from the PVD allowed us to visualize patterns of insula development across the cortex for each insula region. We next sought to annotate the primary axis of insula development by comparing PC1 scores to known patterns of brain organization. We compared the primary and secondary axis of insulo-cortical development (primary and secondary developmental scores from first PCA) of all edges to both the sensorimotor-association axis (S-A axis) and known canonical networks.

#### Sensorimotor-Association Axis

The S-A axis was derived from multi-modal brain maps that exhibit stereotyped feature variability between primary sensorimotor and transmodal association cortices. The S-A axis assignments used for this study were parcellated using the same atlas as above (HCP-MMP atlas). We used Spearman’s rank correlations to quantify associations between observed PC1 scores for each insula region and S-A axis ranks (Sydnor et al., 2021, publicly available and obtained from http://github.com/PennLINC/S-A_ArchetypalAxis). PC1 scores for the insula regions were averaged across hemispheres. Statistical significance between the PC1 and S-A axis was tested using spin-based spatial permutation tests as described previously^110^. A *p* value (*p*spin) was computed by comparing the empirically observed correlation to a null distribution of correlations from 1000 spins. A statistically significant result indicated alignment of an insula region’s age-related trajectories with the S-A axis.

#### Network enrichment

We next tested whether age-related trajectories (PC1 scores) were enriched within canonical cortical networks. Cortical regions were assigned to 12 functional networks using the Cole-Anticevic Brain-wide Network Partition (CAB-NP)^56^. Given the large number of edges, we focused on the bilateral regions from the anterior and posterior poles: the anteroventral and agranular regions at the anterior pole and the granular insula at the posterior pole. In accordance with the antero-posterior organization of the insula, we surmised that these regions would likely demonstrate the greatest difference in developmental pattern. For each of the 12 functional networks, we calculated the empirical mean PC1 score across assigned cortical regions. This resulted in 12 mean PC1 values – one corresponding to each network for each insula region.

Statistical significance of mean network patterns was assessed using a modified spin test, where a null distribution of network means was generated by computing mean PC1 scores after 1000 spatially rotated network assignments. Two-tailed *p*-values were calculated as the proportion of permutations with an absolute deviation from the null distribution greater than the observed statistic. A statistically significant result would indicate that the primary axis of development (i.e., PC1) of an insula region is preferentially expressed within certain functional networks.

### Characterizing differences in developmental patterns across the insula

We next sought to annotate the results from the second step of the PVD, which evaluated how different insula regions align with dominant age-related trajectories. First, we extracted the second PCA scores, representing variance in the developmental connectivity profiles of the insula regions. We also sought to understand the overall cortical pattern of development. We hypothesized that the insula would develop across the S-A axis, so we tested cortical loadings from the second PCA to the S-A axis. As above, statistical significance between the PC1 loadings and S-A axis was tested using spin-based spatial randomization tests^110^. A *p* value (*p*spin) was computed by comparing the empirically observed correlation to a null distribution of correlations from 1000 spins. A statistically significant result indicated alignment of overall insulo-cortical age-related trajectories with the S-A axis.

### Vertex-Level Analysis

Our main parcel-wise analyses provided strong evidence that different regions of the insula have divergent patterns of functional connectivity development. As a final step, we sought to relate these patterns to known gradients in insular cytoarchitecture. To do so, we conducted a high-resolution vertex-wise analysis – the HCP-MMP atlas was simply too coarse to capture fine-grained gradients of insula functional connectivity development and cytoarchitecture.

#### Vertexwise polynomial regression

The parcel-wise analysis using GAMs of the HCP-MMP atlas revealed >99% of the variance could be explained by two components – one primarily linear and one primarily quadratic. Given that the developmental patterns of the vertex-level data is expected to closely follow that of the ROI analysis, we conducted a vertex-level analysis using a polynomial regression with vertex-wise insulo-cortical functional connectivity as the outcome and age, age^2^, sex and in-scanner motion as independent variables. As above, we summarized in-scanner head motion as mean FD averaged across all runs for each participant. Age was *z*-scored prior to generating the quadratic term to reduce collinearity. We extracted linear and quadratic estimated regression coefficients for each edge. To allow for more direct comparison of age- and age^2^-related effects, we normalized each age and age^2^ coefficient by dividing by the standard deviation of all age and age^2^ terms, respectively.

#### Relating insulo-cortical functional connectivity development to insula microstructure

As with our main parcel-wise analysis, relating insulo-cortical functional connectivity development to cortical properties – such as insula cytoarchitecture – required data reduction. Similar to our prior analyses, PCA (*prcomp* from *stats* package version 4.4.1 in R) was performed to identify patterns of insulo-cortical development based on the vertex-level analysis. Specifically, PCA was applied to a matrix with both the normalized linear and quadratic coefficients. In this matrix, insula vertices were rows while columns contained linear and quadratic coefficients from cortical vertices. We used the BigBrainWarp toolbox ^114^ to examine the association between microstructural profiles of the insula to developmental patterns captured by PCA (PC1 scores). The BigBrainWarp toolbox provides a microstructural atlas derived from MRI-based intracortical intensity profiles sampled across multiple equidistant cortical surfaces. These profiles were then used to compute an inter-regional similarity matrix from which first and second gradient scores were derived. These gradients summarize the dominant axes of variation in microstructural architecture across the cortex^115^ derived from multilayer patterns of myelination and corresponding with laminar organization and von Economo classification of cytoarchitecture ^59^. We were unable to use spin tests to test for statistical significance, as our findings were limited to a section of cortex (the insula) and spin tests require the whole cortical surface to appropriately utilize the spherical projection. Therefore, statistical significance was tested using Moran randomization (adapted from Wagner et al 2015)^60^. Moran randomization creates null distributions with similar spatial autocorrelation. A statistically significant result would indicate that the primary axis of development in the insula aligns with microstructural profiles of the insula.

### Analyses detailing insula development with subcortical structures

The insula has known strong and functionally meaningful connections with subcortical regions, which are often overlooked or underpowered in whole-brain connectivity studies. To provide a comprehensive view of insula connectivity development, we also separately evaluated insula developmental patterns with subcortical regions. We used Tian labels^61^, which parcellates the subcortex into 16 regions, and extracted Pearson *r* connectivity values, which were the average timeseries correlation between vertices of each insula and subcortical region. Hemispheres were averaged. As above, GAMs were fit for each insulo-subcortical edge with resting-state connectivity as the dependent variable, age as a smooth term, and both sex and in-scanner motion as linear covariates:

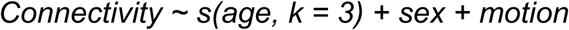

In-scanner head motion was quantified for each participant as mean FD averaged across all included runs. Age was modeled using thin plate regression splines, with the maximum basis complexity (k) set to 3 to reduce overfitting. The effect size of age-related changes for each insulo-subcortical edge was quantified as the partial R^2^ comparing a full model and reduced model without the age term. The direction of change (increasing vs decreasing connectivity with age) was determined from the average derivative (*gratia*^113^, version 0.10.0.900). Significant age effects were identified using ANOVA with false discovery rate (FDR) correction (*Q* = 0.05). To visualize estimated smooth function, we generated predicted values from the fitted GAM for 100 evenly spaced values across the age range using *predict* from mgcv package (Wood, 2025, version 1.9-1). All analyses were conducted with R.

Data availability statement: The present work used raw data from the Human Connectome Project-Development (HCP-D) through data Release 2.0 and Human Connectome Project: Young Adult (HCP-YA) through data Release S1200. Data is available for download through the NIMH Data Archive (https://nda.nih.gov/). Analysis code is available at https://github.com/pennlinc/Flook_insula_dev with a detailed guide to code and procedures for replicating analyses.

## Funding sources

R25MH132513 (EAF), R13DA060048 (EAF), R25MH119043 (EAF/CH), 3R01EB022573 (TDS), 2R01MH112847 (TDS & RTS), 2R01MH113550 (TDS), 2R37MH125829 (TDS)

## Acknowledgments

This publication was made possible through the American Academy of Child and Adolescent Psychiatry (AACAP) Pilot Research Award for Early Career Faculty and Child and Adolescent Psychiatry Fellows, supported by AACAP and KTGF; its contents are the responsibility of the authors and do not necessarily reflect the official views of AACAP and KTGF.

Research reported in this publication was supported by the National Institute of Mental Health of the National Institutes of Health under Award Number U01MH109589 and by funds provided by the McDonnell Center for Systems Neuroscience at Washington University in St. Louis.

Data were provided by the Human Connectome Project, WU-Minn Consortium (HCP-YA 1U54MH091657, Principal Investigators: David Van Essen and Kamil Ugurbil; HCP-D U01MH109589, Principal Investigators: David Van Essen, Susan Bookheimer, Randy L Buckner, Deanna Barch, Mirella Dapretto, Stephen Mark Smith, Leah Helene Somerville, Kathleen M Thomas, and Essa Yacoub) funded by the 16 NIH Institutes and Centers that support the NIH Blueprint for Neuroscience Research; and by the McDonnell Center for Systems Neuroscience at Washington University.

## Disclosures

AAB has equity in, has consulted for, and has an inventorship interest in Children’s Hospital of Philadelphia intellectual property licensed to Centile Bioscience.

## Notes

### Competing Interest Statement

Aaron Alexander-Bloch has equity in, has consulted for, and has an inventorship interest in Children's Hospital of Philadelphia intellectual property licensed to Centile Bioscience.

