## Supplementary Figure 7 for "Insula functional connectivity development follows a canonical antero-posterior axis of organization"

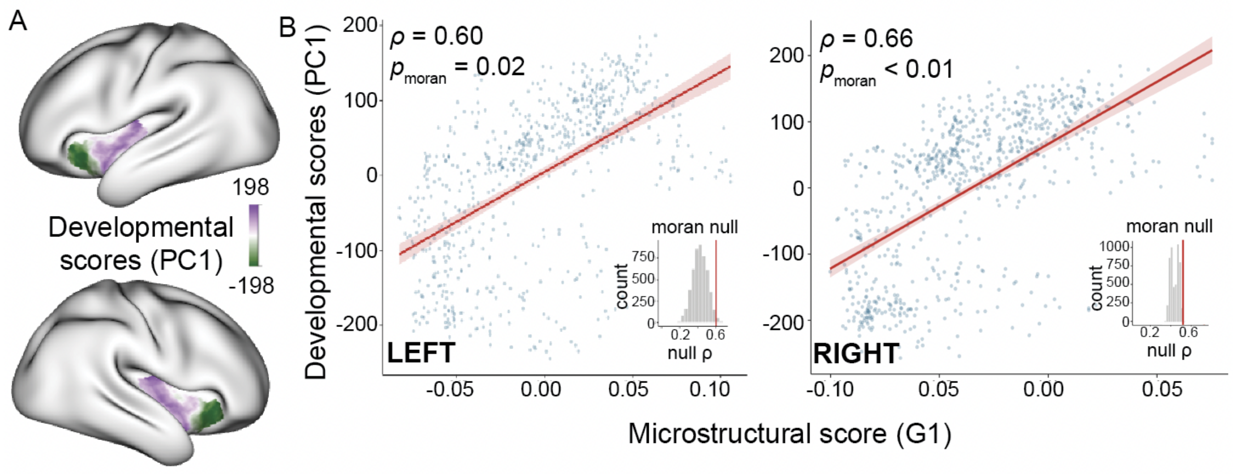


**Supplementary Figure 7:** Developmental gradient of the insula aligns with underlying cytoarchitecture of the insula | (**a**) Vertex-level analyses demonstrate a transition from anterior to posterior insula, reflecting a similar pattern of age-related trajectories as the parcellated approach. (b) Developmental patterns across the insula correlate with underlying microstructure (left insula correlation with primary microstructural gradient shown; $\rho = 0.60$, $p_{Moran} = 0.02$, right insula correlation with primary microstructural gradient shown: $\rho = 0.66$, $p_{Moran} = 0.003$). Microstructure is defined from the BigBrain atlas derived from MRI-based intracortical intensity profiles; these microstructural scores have been shown to be highly correlated with underlying cytoarchitecture. Correlation values determined by Spearman correlation; statistical significance evaluated using Moran randomization.
