## Supplementary Figure 6 for "Insula functional connectivity development follows a canonical antero-posterior axis of organization"

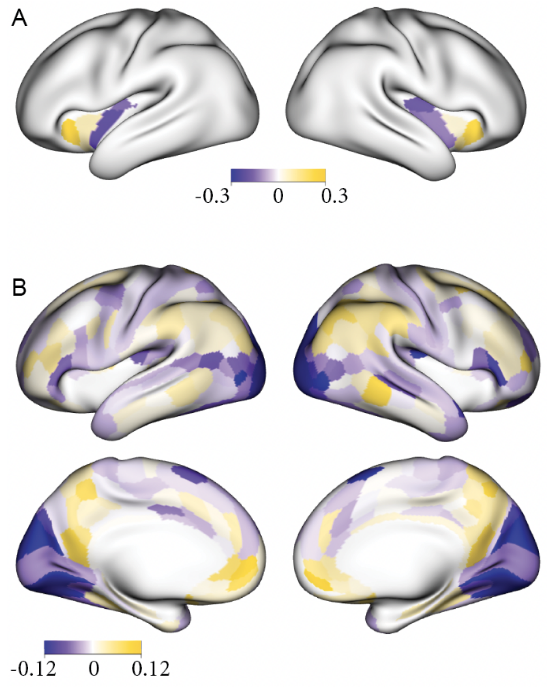


**Supplementary Figure 6:** Summary of the secondary developmental pattern demonstrated regional heterogeneity across the insula and cortex | (**a**) Scores from the second step of the PVD summarize how strongly each insula subregion expresses the secondary pattern of insulo-cortical development. Anterior insula subregions showed similar scores that differed from those of posterior subregions, revealing an antero-posterior organization of developmental trajectories. (**b**) Cortical loadings from the same PVD component show that the cortical regions that most strongly contribute to this secondary developmental pattern.
