## Supplementary Figure 5 for "Insula functional connectivity development follows a canonical antero-posterior axis of organization"

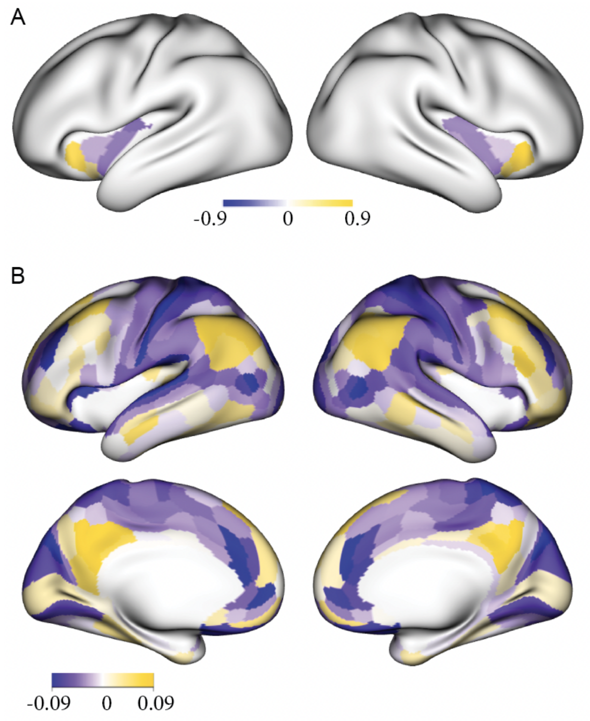


**Supplementary Figure 5:** Summary of the primary developmental pattern demonstrated regional heterogeneity across the insula and cortex | (**a**) Scores from the second step of the PVD summarize how strongly each insula subregion expresses the dominant pattern of insulo-cortical development. Anterior insula subregions showed similar scores that differed from those of posterior subregions, revealing an antero-posterior organization of developmental trajectories. (**b**) Cortical loadings from the same PVD component show that the cortical regions that most strongly contribute to this primary developmental pattern.
