## Supplementary Figure 4 for "Insula functional connectivity development follows a canonical antero-posterior axis of organization"

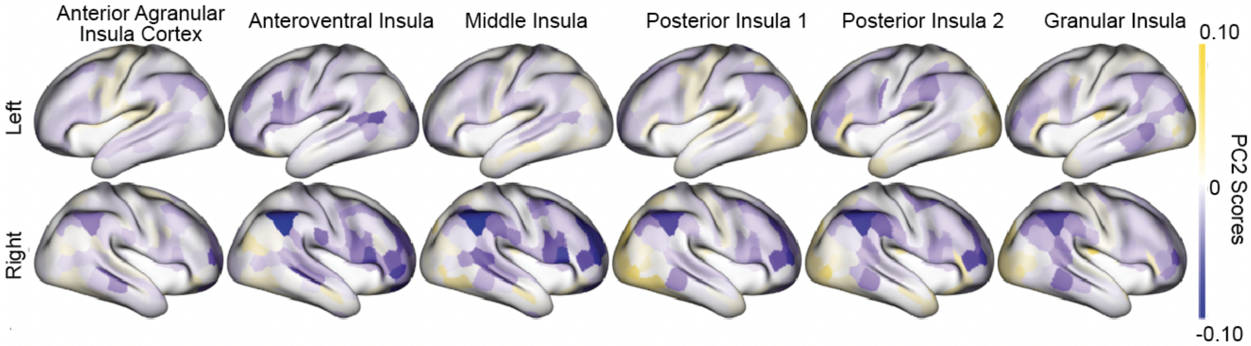


**Supplementary Figure 4:** Cortical patterns of secondary component of insulo-cortical age-related trajectories vary across insula subregions | The cortical distribution of PC2 scores along the secondary developmental axis of insulo-cortical connectivity varied across insula subregions. These values are the extracted PC2 scores from the first step of the PVD, representing the secondary developmental pattern of connectivity across all edges, arranged such that the individual insula regions are visualized separately.
