## Supplementary Figure 3 for "Insula functional connectivity development follows a canonical antero-posterior axis of organization"

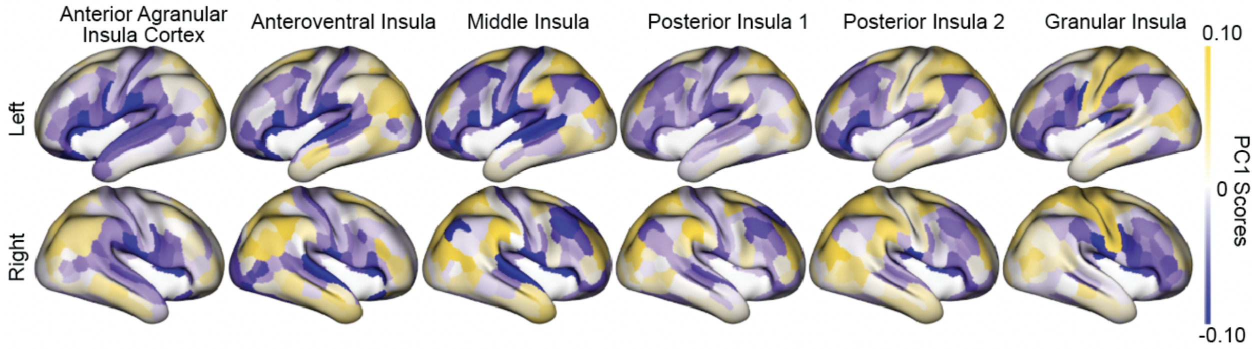


**Supplementary Figure 3**: Cortical patterns of primary component insulo-cortical age-related trajectories vary across insula subregions | The cortical distribution of PC1 scores along the primary developmental axis of insulo-cortical connectivity varied across insula subregions. These values are the extracted PC1 scores from the first step of the population value decomposition (PVD), representing the primary developmental pattern of connectivity across all edges, arranged such that the individual insula regions are visualized separately. Positive scores reflect greater connectivity in older ages; negative scores are lesser connectivity in older age.
