## Supplementary Figure 2 for "Insula functional connectivity development follows a canonical antero-posterior axis of organization"

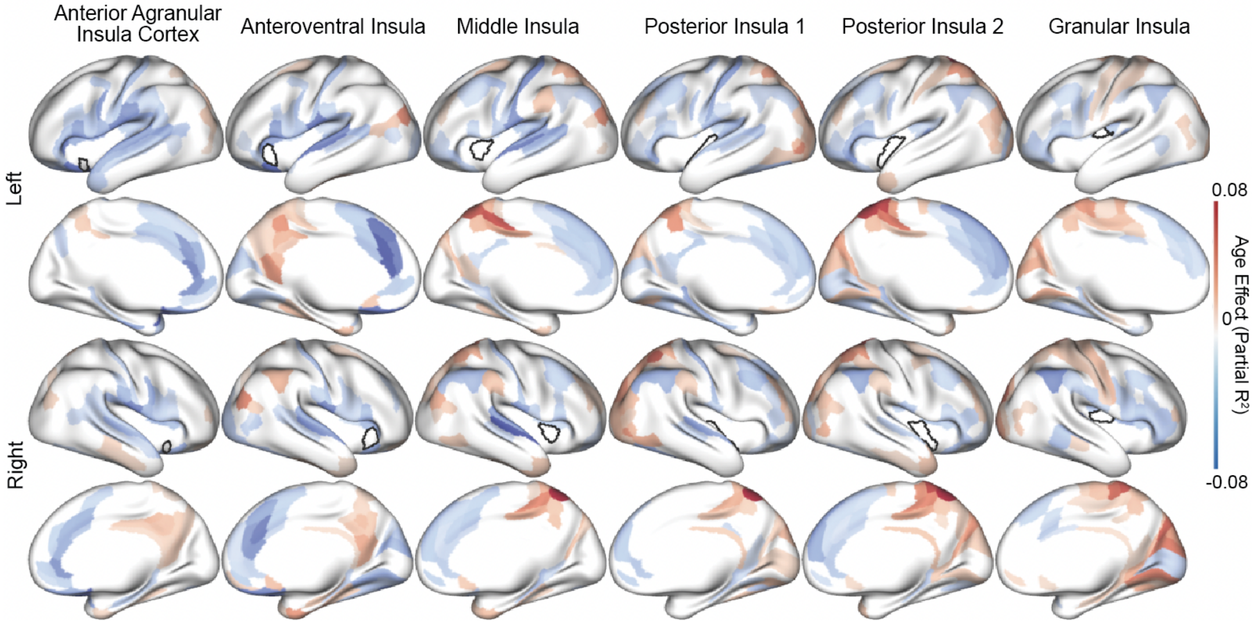


**Supplementary Figure 2:** Effect of age on insulo-cortical connectivity varies across insula subregions | Cortical distribution of age effects was quantified using partial $R^2$, calculated by comparing Generalized Additive Models (GAMs) with and without an age term. Multiple comparisons were controlled using the false discovery rate (FDR). Colors reflect the direction and strength of significant age effects (positive = red, negative = blue), significance defined as $p_{FDR} < 0.05$, with non-significant cortical subregions shown in white.
