## Supplementary Figure 1 for "Insula functional connectivity development follows a canonical antero-posterior axis of organization"

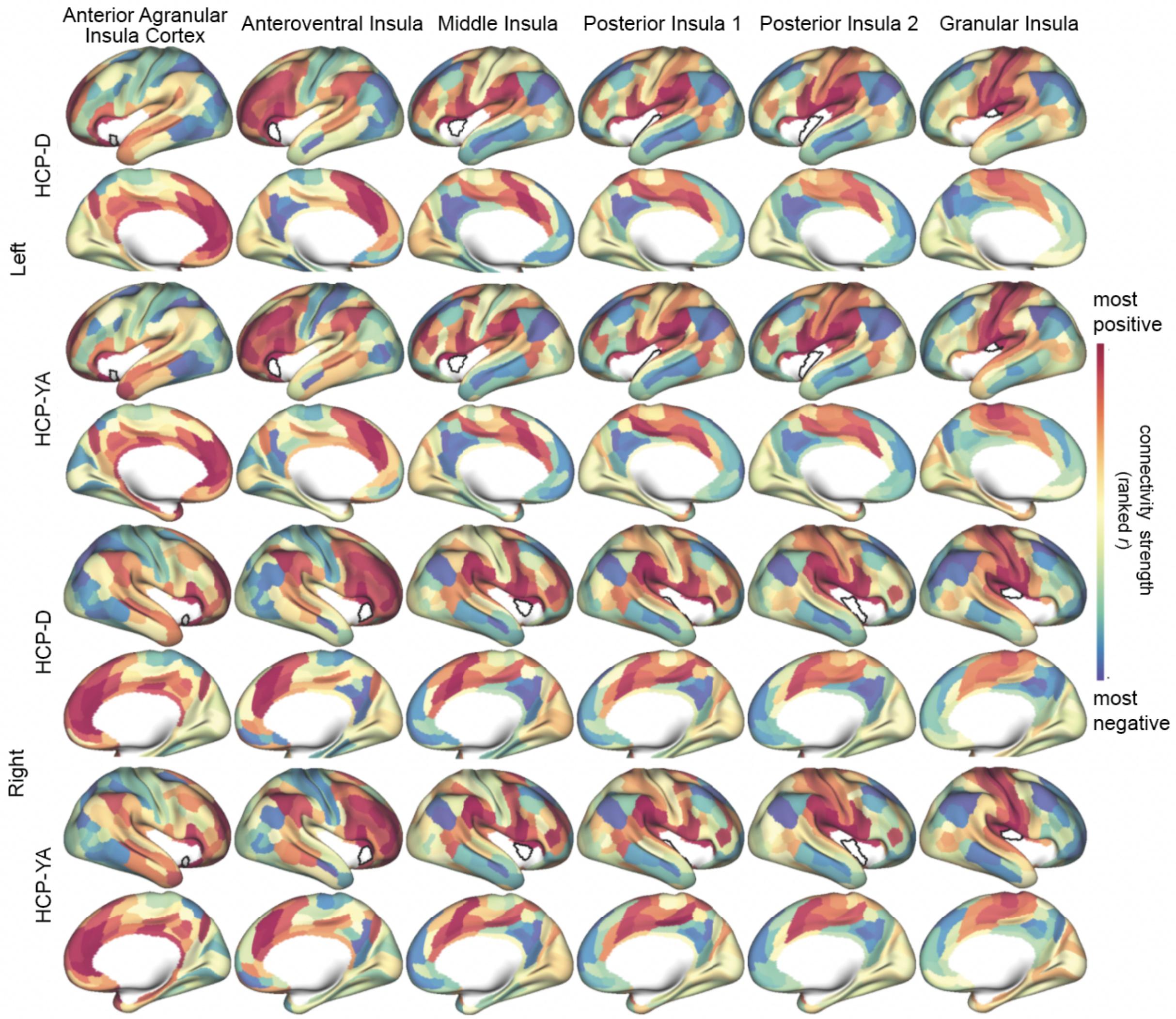


**Supplementary Figure 1:** Connectivity with the cortex varies across insula subregions and is globally similar between youth and adulthood | Cortical surface maps rank order of insulo-cortical connectivity between HCP-D and HCP-YA for each insula subregion (one subregion per column). Subregions are organized from most anteroventral on the left (anterior agranular insula cortex) to most posterodorsal on the right (granular insula). Correlations are shown in rank order from strongest positive (red) to strongest negative (blue) connectivity, with minimal connectivity in the middle (yellow). Correlation between HCP-YA and HCP-D connectivity maps for all subregions > 0.91, FDR $p_{spin}$ < 0.001. Black outlines represent the location of the insula subregions.
