## Supplementary Table 2 for "Insula functional connectivity development follows a canonical antero-posterior axis of organization"

**Supplementary Table 2:** Insulo-subcortical connectivity analyses revealed few age-related changes. We used generalized additive models to capture the effect of age on connectivity between insula subregions and subcortical structures. Age effects were quantified as the partial $R^2$ comparing a model with and without the age term. AAIC: anterior agranular insula cortex; AVI: anteroventral insula; Ig: granular insula; MI: middle insula; PoI1: posterior insula 1; PoI2: posterior insula 2; AMY: amygdala; CAU: caudate; GP: globus pallidus; HIP: hippocampus; NAc: nucleus accumbens; PUT: putamen; THAL_ant: anterior thalamus; THAL_post: posterior thalamus

| Insula subregion | Subcortical region | ANOVA p-value | FDR corrected p-value |
|---|---|---|---|
| PoI2 | HIP | 0.440 | 0.820 |
| MI | HIP | 0.180 | 0.730 |
| AVI | HIP | 0.790 | 0.870 |
| AAIC | HIP | 0.660 | 0.820 |
| PoI1 | HIP | 0.110 | 0.730 |
| Ig | HIP | 0.044 | 0.520 |
| PoI2 | THAL_post | 0.200 | 0.730 |
| MI | THAL_post | 0.560 | 0.820 |
| AVI | THAL_post | 0.820 | 0.870 |
| AAIC | THAL_post | 0.170 | 0.730 |
| PoI1 | THAL_post | 0.230 | 0.730 |
| Ig | THAL_post | 0.640 | 0.820 |
| PoI2 | THAL_ant | 0.310 | 0.820 |
| MI | THAL_ant | 0.850 | 0.890 |
| AVI | THAL_ant | 0.580 | 0.820 |
| AAIC | THAL_ant | 0.620 | 0.820 |
| PoI1 | THAL_ant | 0.290 | 0.820 |
| Ig | THAL_ant | 0.120 | 0.730 |
| PoI2 | PUT | 0.550 | 0.820 |
| MI | PUT | 0.670 | 0.820 |
| AVI | PUT | 0.350 | 0.820 |
| AAIC | PUT | 0.220 | 0.730 |
| PoI1 | PUT | 0.420 | 0.820 |
| Ig | PUT | 0.660 | 0.820 |
| PoI2 | CAU | 0.870 | 0.890 |
| MI | CAU | 0.730 | 0.850 |
| AVI | CAU | 0.810 | 0.870 |
| AAIC | CAU | 0.550 | 0.820 |
| PoI1 | CAU | 0.780 | 0.870 |
| Ig | CAU | 0.210 | 0.730 |
| PoI2 | AMY | 0.420 | 0.820 |
| MI | AMY | 0.120 | 0.730 |
| AVI | AMY | 0.490 | 0.820 |
| AAIC | AMY | 0.680 | 0.820 |
| PoI1 | AMY | 0.230 | 0.730 |
| Ig | AMY | 0.460 | 0.820 |
| PoI2 | NAc | 0.520 | 0.820 |
| MI | NAc | 0.940 | 0.940 |
| AVI | NAc | 0.600 | 0.820 |
| AAIC | NAc | 0.330 | 0.820 |
| PoI1 | NAc | 0.480 | 0.820 |
| Ig | NAc | 0.440 | 0.820 |
| PoI2 | GP | 0.011 | 0.270 |
| MI | GP | 0.001 | 0.037 |
| AVI | GP | 0.032 | 0.520 |
| AAIC | GP | 0.480 | 0.820 |
| PoI1 | GP | 0.080 | 0.730 |
| Ig | GP | 0.340 | 0.820 |
