## Supplementary Table 1 for "Insula functional connectivity development follows a canonical antero-posterior axis of organization"

**Supplementary Table 1**: Network enrichment was quantified by averaging first principal component (PC1) scores for the anterior agranular insula cortex (AAIC), anteroventral insula (AVI), and granular insula (Ig) within each cortical network. Cortical networks were assigned using the Cole atlas[1]. Statistical significance was assessed using spin-based spatial permutation testing, in which network labels were spatially rotated to generate null distributions of mean PC1 scores while preserving spatial autocorrelation. Multiple comparisons were controlled using the Benjamini-Hochberg false discovery rate (FDR) correction across all insula subregion–network comparisons.

| Insula subregion | Cortical network | Observed mean PC1 Score | Spin test p-value | FDR-corrected p-value |
|---|---|---|---|---|
| AAIC | Auditory | −0.042 | 0.506 | 0.772 |
| AAIC | Cingulo-Opercular | −0.051 | 0.006 | 0.108 |
| AAIC | Default | −0.005 | 0.708 | 0.772 |
| AAIC | Dorsal-Attention | 0.017 | 0.137 | 0.385 |
| AAIC | Frontoparietal | 0.008 | 0.166 | 0.427 |
| AAIC | Language | −0.052 | 0.035 | 0.222 |
| AAIC | Orbito-Affective | −0.093 | 0.037 | 0.222 |
| AAIC | Posterior-Multimodal | 0.003 | 0.675 | 0.772 |
| AAIC | Somatomotor | −0.027 | 0.647 | 0.772 |
| AAIC | Ventral-Multimodal | 0.009 | 0.659 | 0.772 |
| AAIC | Visual1 | 0.017 | 0.466 | 0.763 |
| AAIC | Visual2 | 0.016 | 0.322 | 0.644 |
| Ig | Auditory | −0.055 | 0.137 | 0.385 |
| Ig | Cingulo-Opercular | −0.008 | 0.343 | 0.644 |
| Ig | Default | 0.005 | 0.772 | 0.816 |
| Ig | Dorsal-Attention | 0.011 | 0.957 | 0.957 |
| Ig | Frontoparietal | −0.005 | 0.358 | 0.644 |
| Ig | Language | −0.017 | 0.229 | 0.545 |
| Ig | Orbito-Affective | −0.006 | 0.675 | 0.772 |
| Ig | Posterior-Multimodal | 0.065 | 0.099 | 0.385 |
| Ig | Somatomotor | 0.050 | 0.139 | 0.385 |
| Ig | Ventral-Multimodal | −0.002 | 0.794 | 0.816 |
| Ig | Visual1 | 0.028 | 0.647 | 0.772 |
| Ig | Visual2 | 0.042 | 0.304 | 0.644 |
| AVI | Auditory | −0.078 | 0.138 | 0.385 |
| AVI | Cingulo-Opercular | −0.051 | 0.010 | 0.120 |
| AVI | Default | 0.025 | 0.032 | 0.222 |
| AVI | Dorsal-Attention | 0.056 | 0.001 | 0.036 |
| AVI | Frontoparietal | −0.002 | 0.634 | 0.772 |
| AVI | Language | −0.047 | 0.088 | 0.385 |
| AVI | Orbito-Affective | −0.055 | 0.406 | 0.696 |
| AVI | Posterior-Multimodal | 0.059 | 0.075 | 0.385 |
| AVI | Somatomotor | −0.023 | 0.687 | 0.772 |
| AVI | Ventral-Multimodal | 0.059 | 0.242 | 0.545 |
| AVI | Visual1 | 0.016 | 0.584 | 0.772 |
| AVI | Visual2 | −0.025 | 0.610 | 0.772 |
